# Single-Cell Profiling of Dynamic Epicardial Cell States During Myocardial Infarction

**DOI:** 10.64898/2026.08.31.748312

**Authors:** David Wong, Jenny Cheng, Xia Yang, Pearl Quijada

## Abstract

**Background:** The epicardium is reactivated after myocardial infarction (MI); however, the gene expression profiles of post-MI adult epicardial subpopulations remain incompletely defined.

**Methods:** Single-cell RNA sequencing was performed on lineage-traced *Wt1*^+^ epicardial cells from *Wt1^CreERT^*^2^*^/+^; R26^tdT/+^; Pdgfra^nGFP/+^* adult mice after sham surgery or at 7 and 14 days after permanent artery ligation to induce MI. Immunostaining was performed on *Wt1*-lineage-traced cardiac tissue to validate spatial expression after ischemic injury.

**Results:** Unbiased clustering identified nine transcriptionally distinct epicardial populations, encompassing mesothelial, fibroblast/mesenchymal, transitional, and proliferative phenotypes. Fibroblast-like epicardial cells (Wt1^+^/Pdgfra^+^) showed time-dependent expression profiles associated with upregulation of epithelial-to-mesenchymal transition (EMT) and extracellular matrix (ECM) gene programs. At 7 days post-MI, there was notable enrichment of genes related to chemokines and Wnt components. By 14 days post-MI, the expression profile shifted toward immune regulation. In contrast, a Wt1^high^/Msln^+^ population showed minimal upregulation of EMT gene programs but enhanced paracrine signaling related to wound healing and semaphorins, suggesting reactivation of reparative and angiogenic functions akin to those of the epicardium during embryonic development. Immunostaining and in situ hybridization fluorescence analyses validated laminar epicardial cell placement after MI, comprising a surface *Msln*^+^ sheet, an overlapping *Wt1*-lineage band, and a subadjacent PDGFRα^+^ and Periostin^+^ compartment that expands 7-14 days after MI and regresses by day 28 post-ischemia.

**Conclusions:** Our data define epicardial gene programs in which a signaling epithelial cell surface overlays an effector mesenchymal cell stroma to coordinate angiogenesis, leukocyte recruitment, and ECM remodeling. This study presents the first integrated single-cell atlas of epicardial-derived cells across multiple post-ischemic timepoints, offering new insights into their reparative potential and dynamic signaling diversity in the injured adult heart.

## INTRODUCTION

During murine embryonic cardiac development, specifically between embryonic days (E) 9.5 and 11.5, the epicardium envelops the heart and plays a critical role in secreting paracrine factors that facilitate the growth and maturation of the myocardium and coronary vasculature^1^. By approximately E12.5, the epicardium undergoes epithelial-to-mesenchymal transition (EMT), a process that generates epicardium-derived progenitor cells capable of differentiating into fibroblasts, smooth muscle cells, and pericytes, and provides indispensable extracellular matrix (ECM) to the rapidly expanding myocardium^2^. Recent single-cell transcriptional studies have highlighted the epicardium’s dynamic capacity to act as a non-cell-autonomous regulator, orchestrating embryonic angiogenesis and the maturation of the coronary vasculature and myocardium through a specific temporal-spatial code of paracrine signaling. Comparative studies in adult zebrafish and salamanders, along with early neonatal mammalian models, have highlighted the epicardium as a central driver of robust cardiac regeneration, in part by reactivating developmental epicardial programs after injury^3–5^. This regenerative process is mediated by paracrine factors, including Fibroblast Growth Factors (FGFs), Insulin-like Growth Factors (IGFs), Platelet-Derived Growth Factors (PDGFs), as well as Wnt and retinoic acid (RA) signaling^6^. To advance therapeutic strategies for repairing the injured adult heart, it is crucial to harness and modulate gene programs from structures such as the epicardium, which exhibit remarkable reparative abilities, given the limited self-repair capacity of adult mammalian hearts.

In the adult heart, the epicardium is quiescent under homeostatic conditions. However, following ischemic injury, epicardial cells are re-activated and contribute to several key reparative processes, including the modulation of inflammation^7^, deposition of ECM^8,9^, recruitment of mural cells^10,11^, and stimulation of angiogenesis^4^, in part through induction of EMT and the formation of a multilayered epicardial stromal cell compartment at the heart surface. In the infarcted murine heart, epicardial cells proliferate at the heart surface and upregulate embryonic genes, including *Wt1*, *Tbx18*, and *Raldh2*^4^, as well as developmental signaling pathways such as Wnt^12^ and TGFβ^13^. While it is generally expected that the adult post-injury epicardium recapitulates embryonic functions, emerging evidence suggests distinct molecular signaling pathways between the embryonic and adult epicardium^14^. Single-cell comparisons of fetal and aged adult epicardial cells, as well as adult murine post-MI epicardial cells, demonstrate that the aged human epicardium, as well as the post-MI epicardium, contains more fibrotic and immune-modulatory cellular subsets and lacks key pro-regenerative angiogenic programs characteristic of the fetal epicardium^9,14^. Nonetheless, adult epicardial cells have been shown to secrete various reparative factors that promote endothelial cell activation^15^, neovascularization^16,17^, and cardiomyocyte survival^18^ through paracrine signaling. As such, the adult epicardium maintains its status as an important cellular reservoir and a vital signaling hub with cardiogenic potential in the setting of an injured adult heart, and represents a potential therapeutic target for reprogramming in ischemic heart disease.

Despite the epicardium’s various roles during cardiac repair, there is a paucity of studies investigating the cellular and molecular heterogeneity of the adult epicardium, particularly across distinct temporal phases of ischemia. Previous studies examining the epicardium have been limited by the inclusion of contaminating stromal cell types at a single time point after ischemic injury or by reliance on bulk, non-lineage-tracing transcriptomic approaches^9,19,20^, which make it challenging to distinguish bona fide epicardial-derived cells from other interstitial mesenchymal cells. As a result, understanding how distinct epicardial subpopulations emerge and function during the dynamic phases of cardiac repair remains incomplete. To address this, we performed single-cell RNA sequencing (scRNA-seq) of epicardial cells isolated from sham, 7-day, and 14-day post-MI hearts, representing key stages of the ischemic cardiac repair process^21^. To lineage trace epicardial cells, we utilized a triple transgenic mouse model, *Wt1^CreERT^*^2^*^/+^*; *R26^tdT/+^*; *Pdgfra^nGFP/+^*, which enables fluorescence-based genetic labeling of *Wt1*-expressing epicardial-derived cells and simultaneous labeling of *Pdgfra*^+^ fibroblasts, which epicardial cells have been reported to transition towards during ischemic injury^22,23^. This model also allows the capture of both mesothelial-like epicardial cells and epicardial-derived fibroblast-like populations within a single-lineage-traced framework. Our approach reveals the dynamic transcriptional programs, cellular heterogeneity, and reparative roles of epicardial-derived populations in the infarcted heart, defines mesothelial, transitional, fibroblast-like, proliferative, and immune-interfacing epicardial states, and identifies stage-specific paracrine programs, including semaphorins, chemokines, and Wnt signaling.

## METHODS

### Experimental animals

All animal studies, including husbandry, breeding, and experimental procedures, were conducted in accordance with protocols approved by the University Committee on Animal Resources at the University of California, Los Angeles, and in accordance with the guidelines of the NIH Guide for the Care and Use of Laboratory Animals. The C57BL/6J mice strain (stock #000664) was obtained from the Jackson Laboratory. A C57BL/6J colony was maintained in our vivarium by crossing C57BL/6J male mice to C57BL/6J female mice. Female mice of the C57BL/6J stock mouse strain from the Jackson Laboratory were introduced into our in-house colony every 10 generations to prevent genetic drift. Wt1-CreERT2 heterozygous males were purchased from the Jackson Laboratory (stock #010912) and crossed to R26-tdTomato homozygous reporter females (stock #007909) to generate heterozygous Wt1-CreERT2; R26-tdTomato mice. Mice expressing the H2B-eGFP fusion gene from the endogenous *Pdgfra* locus (*PDGFRα^nGFP^* heterozygous mice) were purchased from the Jackson Laboratory (stock #007669). All mice were maintained on a C57BL/6J background in a temperature-controlled environment with 12-hour light/dark cycles and received food and water *ad libitum* in either cages of a maximum of 5 females or 4 males per cage.

Detailed descriptions of all experimental procedures and materials are available in the Supplemental Material. Single-cell RNA sequencing data have been deposited in the National Center for Biotechnology Information Gene Expression Omnibus (GEO) under accession #GSE336312. Fetal epicardial cell single-cell RNA-sequencing can be found under the GEO accession #GSE154715.

## RESULTS

### Confirmation of lineage traced epicardial cells isolated from the adult mouse heart

Prior studies have investigated adult epicardial cell function in the contexts of human aging and following short-term ischemic injury in murine models^9,14^. Yet, a comprehensive timeline outlining the transcriptional profile of epicardial cells during the transition from the acute to chronic phases of MI remains unclear. To identify and label adult epicardial cells, we used the *Wt1* tamoxifen-inducible CreERT2 mouse model crossed to a *R26-tdTomato* mouse reporter (*Wt1^CreERT2/+^*; *R26^tdT/+^*), which allows for robust labeling of the homeostatic adult epicardium following tamoxifen administration^4^. To track if epicardial cells expressed quiescent cardiac fibroblast markers, we crossed *Wt1^CreERT2/+^*; *R26^tdT/+^* to the platelet-derived growth factor-α reporter strain (*Pdgfra^nGFP/+^*)^24^, generating a triple transgenic *Wt1^CreERT2/+^*; *R26^tdT/+^*; *Pdgfra^nGFP/+^* mouse that lineage traces *Wt1* cells and co-labels epicardial cells with nuclear GFP upon expression of PDGFRα **(Figure 1A)**. Eight-week-old mice received tamoxifen for 5 days, followed by a 7-day washout period, before undergoing sham or permanent ligation of the left anterior artery to induce MI. Adult hearts were harvested following sham surgery (N=5) and at 7 (N=5) and 14 (N=2) days post-MI **(Figure 1B)**.

**Figure 1.**
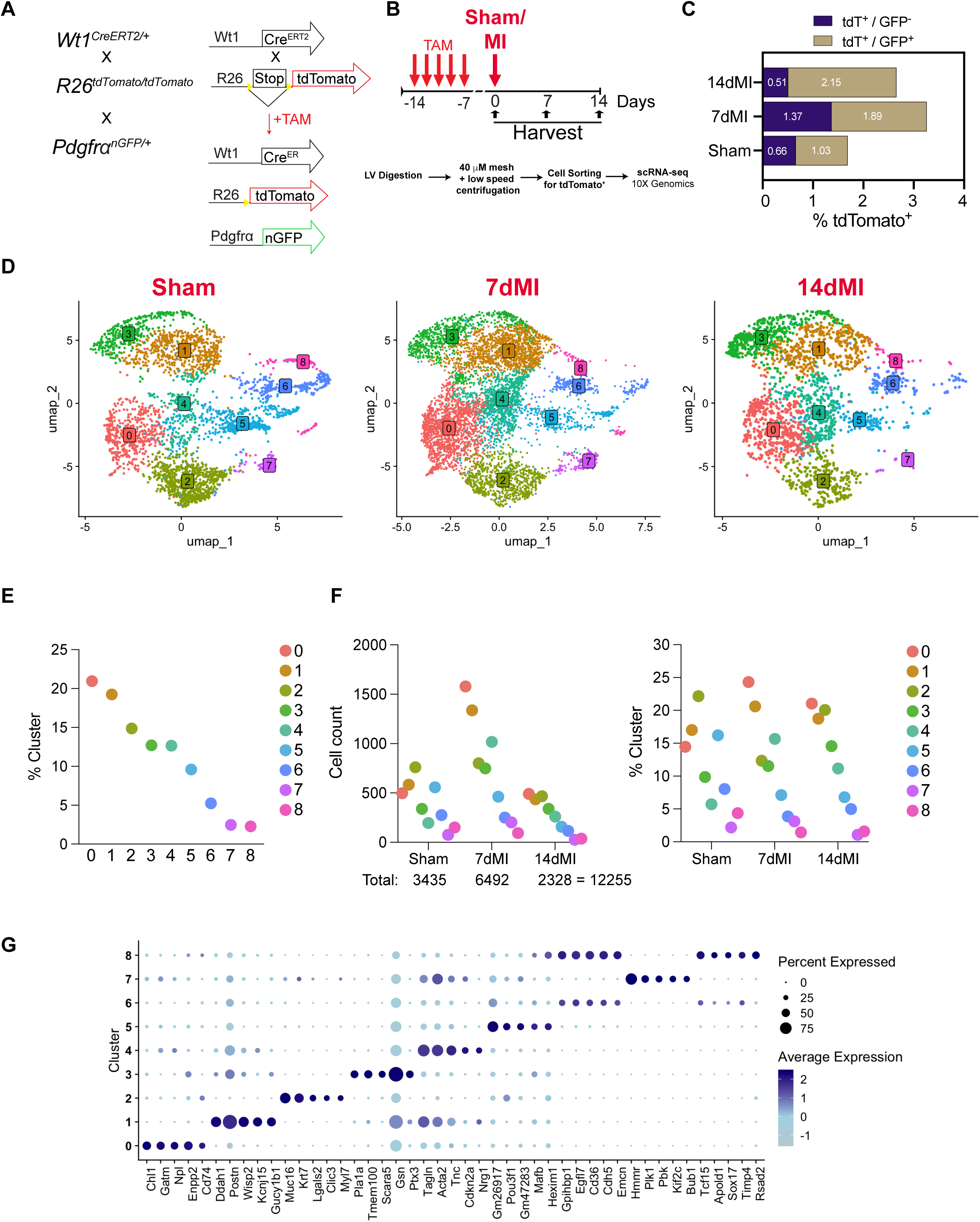
Isolation and analysis of epicardial cells by single-cell RNA sequencing. **(A)** Diagram illustrating the genetic lineage-tracing strategy. *Wt1^CreERT2/+^* mice were crossed with *R26^tdTomato/tdTomato^* mice to mark adult epicardial cells with tdTomato fluorescence (tdT)following tamoxifen (TAM) administration. *Wt1^CreERT2/+^; R26^tdTomato/+^* mice were further crossed to *Pdgfra^nGFP/+^* reporter mouse strain to mark resident adult fibroblasts. **(B)** A diagram illustrating the workflow and timeline of the experiment. TAM was administered two weeks before surgery for five consecutive days, followed by a week washout period, before performing sham or myocardial infarction (MI) surgeries. Hearts were extracted from sham-operated mice and from mice subjected to MI for 7 and 14 days. Left ventricles (LVs) were microdissected, enzymatically digested, filtered, and tdTomato^+^ cells were purified by fluorescence-activated cell sorting (FACS) before performing single-cell RNA sequencing using the 10x Chromium platform. **(C)** Representative percentage of tdTomato^+^ cells recovered across sham, 7, and 14-day MI conditions following LV digestion and FACS. The percentage of cells was subdivided into tdTomato^+^/GFP^-^ and tdTomato^+^/GFP^+^ fractions. Data is represented as one biological sample per condition. **(D)** UMAP visualization of epicardial cell clusters (0-8) in Sham, 7dMI, and 14dMI conditions. **(E)** The cluster percentage across all conditions. **(F)** Cell counts and cluster percentage in sham, 7dMI, and 14dMI conditions. **(G)** Dot plot of the top 5 marker genes across clusters 0-8. Data are presented as biological replicates: N=5 for Sham, N=5 for 7dMI, and N=2 for 14dMI.

Injury time points selected were to capture peak epicardial activation, characterized by increased epicardial thickening and maximal re-expression of fetal epicardial genes at 7 days post-MI (7dMI), followed by the resolution phase, with downregulation of developmental genes at 14 days post-MI (14dMI)^4^. Hearts from each surgery group (Sham and MI) were combined, enzymatically dissociated, and filtered before fluorescent-activated cell sorting (FACS) of all viable tdT^+^ epicardial lineage cells, regardless of GFP positivity **(Figure 1B and Supplemental Figures 1A-C)**.

Representative FACS plots show the expansion of tdT^+^ epicardial-derived cells after MI, with both tdT^+^ and tdT^+^/GFP^+^ populations increasing relative to sham, an indicator of epicardial activation and increased fibroblast-like conversion following injury **(Figure 1C and Supplemental Figures 1A-C)**. To validate the cell population following FACS, sorted fractions from uninjured hearts were analyzed for marker gene expression. Canonical epicardial genes (*Wt1*, *Msln*) were elevated in tdT^+^ cells, with or without co-expression of GFP, while fibroblast-associated genes (*Tcf21*, *Col1a1*, *Pdgfra*, *Postn*) were significantly upregulated in GFP^+^ and double-labeled fractions **(Supplemental Figures 2A-F)**. Notably, the double-positive tdTomato^+^/GFP^+^ subset exhibited lower levels of *Tcf21*, *Pdgfra,* and *Postn*, and comparable levels of *Col1a1* compared to the GFP-alone expressing population **(Supplemental Figures 2C-F)**. These data confirm the successful capture of adult epicardial cells and validate gene expression profiles in both *Wt1*-lineage and *Pdgfra*-expressing cell subpopulations.

### Single-cell RNA sequencing analysis reveals diverse subpopulations of adult epicardial cells

Next, FACS-purified epicardial cells were subjected to scRNA-seq using the 10X Genomics Chromium platform, with independent pooled libraries generated per condition/run (sham: N=5 hearts; 7dMI: N=5 hearts; 14dMI: N=2 hearts). To mitigate technical batch effects across sequencing runs, datasets were integrated using Seurat’s SCTransform anchor-based workflow. Following integration, cells from the independent sham and 7dMI runs showed substantial overlap in low-dimensional space and contributed to the same major clusters, supporting minimal run-to-run deviation and reproducible identification of the major epicardial states. Because a single pooled library represented the 14dMI group, reproducibility for that condition could not be assessed to the same extent across independent runs. After quality control filtering **(Supplemental Figures 3A-E)** and SCTransform-based integration across runs, we identified 9 epicardial cell clusters, comprising 3,435 sham cells, 6,492 cells from 7 days post-MI (7dMI), and 2,328 cells from 14 days post-MI (14dMI) **(Supplemental Figures 4A, 4B)**. Integrated UMAP visualization, run-split cluster mapping, cluster composition analysis, and pseudobulk correlation analysis together indicated minimal residual batch effects and broadly reproducible recovery of major epicardial cell states across independent sham and 7dMI sequencing runs **(Supplemental Figure 4D-G)**. Condition-split UMAPs revealed condition-associated redistribution of cells within the integrated transcriptional space, consistent with injury-associated shifts in epicardial cell states **(Figure 1D and Supplemental Figure 4C)**.

Because UMAP visualization is not quantitative for relative abundance, differences in cluster representation across conditions were evaluated separately using cluster proportion analysis **(Figure 1E, 1F)**. Analysis of cluster proportions revealed expansion of clusters 0, 1, 3, and 4 following MI, whereas clusters 2, 5, 6, and 8 were enriched in sham hearts **(Figures 1E,1F)**. Notably, epicardial cell clusters 5, 6, and 8 had fewer unique genes per cell than other subsets **(Supplemental Figure 3C-E)**, suggesting a more quiescent or metabolically less active state and supporting a sham heart origin. The marker genes associated with each cell cluster are presented in two visual formats: a dot plot highlighting the top 5 differentially expressed marker genes **(Figure 1G)** and a heatmap showing the top 10 differentially expressed genes **(Supplemental Figure 5A)**. Furthermore, epicardial clusters were compared with a previously published scRNA-seq dataset of the infarcted murine heart^25^, confirming a largely mesenchymal cell transcriptional phenotype **(Supplemental Figure 5B)**. Of note, Cluster 7 displayed the highest expression of G_2_/M phase genes (*Cdk1, Ccnb2, Top2a, Foxm1*) and proliferation-associated genes (*Mki67, Hist1h2ap*), indicating a proliferative cellular subset, albeit a minor subset **(Supplemental Figures 6A-C)**. Altogether, these analyses reveal heterogeneous subclusters of adult-epicardial lineage cells, as well as the expansion or contraction of specific cellular subsets in response to ischemic injury.

### The adult heart retains a mesothelial-like epicardial cell population

To define epicardial cell states across conditions, we evaluated the expression of established epicardial, mesothelial, and fibroblast markers across clusters and post-MI time points. *Wt1* was broadly expressed across all clusters, except cluster 1, and displayed a transient upregulation at 7dMI followed by a decline by 14dMI **(Figure 2A)**, consistent with an acute but blunted reactivation of fetal epicardial programs after injury described in prior studies^4^. A clear mesothelial program, defined by expression of *Upk3b*, *Efemp1*, *Msln*, *Krt19*, *Aldh1a2*, and *Tbx18*, was significantly enriched in cluster 2 **(Figures 2B-G)**. *Upk3b, Efemp1*, *Msln*, and *Krt19* are validated postnatal mesothelial markers, and their expression is preserved on the outermost surface of the heart^9^. *Upk3b* and *Efemp2* are expressed in human epicardial cells regardless of age^14^. Overall, these data highlight cluster 2 as a non-invasive/migratory cellular subtype, which comprises a mesothelial sheet observed in both the healthy and injured heart.

**Figure 2.**
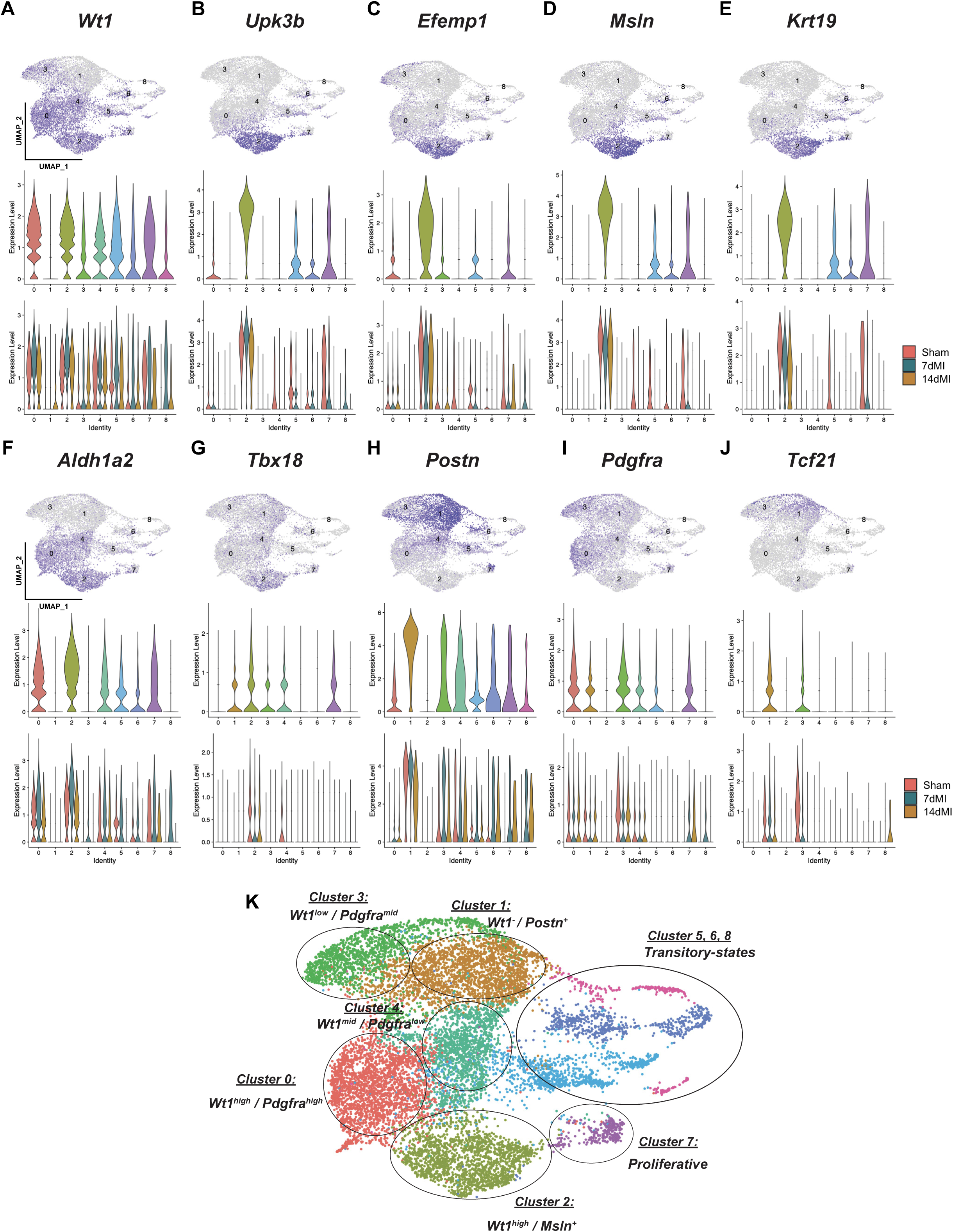
Dynamic expression of gene markers in adult epicardial cells. **(A-J)** Expression of canonical epicardial/mesothelial markers (*Wt1*, *Upk3b, Efemp1, Msln, Krt19, Aldh1a2, Tbx18*) and fibroblast-associated markers (*Postn, Pdgfra, Tcf21*). Top: feature expression (color intensity reflects normalized expression). Middle: violin plots of expression across clusters. Bottom: expression stratified by condition. **(K)** Annotated UMAP summarizing major epicardial states based on marker expression, including: Cluster 0 (*Wt1^high^/Pdgfra^mid^*), Cluster 1 (*Wt1^-^*/*Postn^+^),* Cluster 2 (*Wt1^high^/Msln^+^*), Cluster 3 (*Wt1^low^/Pdgfra^high^*), Cluster 4 (*Wt1^mid^/Pdgfra^low^*), Clusters 5/6/8 (transitory states), and Cluster 7 (proliferative). Data are presented as biological replicates: N=5 for Sham, N=5 for 7dMI, and N=2 for 14dMI.

Cluster 2 represented a significant portion of the total epicardial cells in sham hearts and was reduced in proportion following MI **(Figure 1F)**, as well as in the decline of mesothelial gene expression **(Figures 2B-F)**, suggesting a partial loss or remodeling of the mesothelial layer following the injury. In contrast, *Wt1, Aldh1a2,* and *Tbx18* were more broadly expressed across clusters, supporting the presence of both epithelial-like and stromal epicardial states that retain components of the developmental epicardial program **(Figures 2A, 2F, 2G)**. Additional epicardial cell markers reported in the mouse (*Sfrp2*, *Cd44*, *Pcsk6, Gata5*)^9^ and human epicardium (*Has1, Slpi*, and *Hp*)^14^ were also detected in adult epicardial cells (**Supplemental Figures 7A,7B).** Our data indicate that the adult heart retains a subset of epicardial cells with mesothelial-like characteristics (cluster 2) in both uninjured and MI conditions. Overall, these findings are consistent with transient *Wt1* reactivation following MI; however, *Wt1* combined with preserved *Msln* and *Upk3b* expression indicates a valid mature mesothelial cell population.

### Epicardial cells acquire different degrees of mesenchymal gene signatures following ischemic injury

After MI, epicardial cells have been reported to undergo partial EMT and acquire varying degrees of mesenchymal-like gene signatures following ischemic injury^3,4^. Expression of fibroblast-associated markers exhibited condition- and cluster-specific patterns: *Postn* was most enriched in cluster 1, largely absent from the mesothelial-like cluster 2, and predominantly expressed in post-MI cells across fibroblast-like clusters relative to sham **(Figure 2H)**. *Pdgfra* and *Tcf21* define mesenchymal-like derivatives from the epicardium, with *Tcf21* being essential for the fate specification of epicardial progenitor cells into cardiac fibroblasts during embryonic heart development^1,26^. We found that *Pdgfra* and *Tcf21* expression was largely excluded from the mesothelial cluster 2 and instead enriched in fibroblast-like clusters 0, 1, 3-5, and 7, with clusters 0, 3, and 4 being primarily populated by cells from MI hearts **(Figures 2I, 2J).** *Pdgfra* levels remained relatively stable across sham, 7dMI, and 14dMI, consistent with flow cytometric data indicating that *Wt1*-tdT^+^ captured cells expressed *Pdgfra* before injury, and the number of PDGFRα^+^ cells expanded after MI **(Figures 1C,2I)**. *Tcf21* was restricted to the *Wt1*^-^/*Postn^+^* cluster 1 and *Wt1*^low^ cluster 3, which defines a myofibroblast-like epicardial-derived subset and MI-induced epicardial-derived fibroblasts, respectively **(Figures 2I,2J)**.

Consistent with lineage-tracing studies indicating epicardium-derived cells make minimal contributions towards endothelial lineages during development^27^, we also did not observe expression of endothelial markers (*Cdh5, Efnb2, Kdr, Pecam1, Sox17, Vwf)* within *Wt1*-tdT^+^ cells after MI **(Supplemental Figure 8A)**. We also assessed the expression of pericyte and smooth muscle cell (SMC)-associated genes, as epicardial progenitors contribute to SMC and pericyte lineages during cardiac morphogenesis^1^; however, the extent to which these mural cell types co-express *Wt1* in the adult is not well-defined. Across all *Wt1-*tdT^+^ epicardial-lineage cells, expression of canonical pericyte and SMC markers was largely absent, aside from *Pdgfrb* and *Acta2/Tagln*, which label mature pericytes and SMCs **(Supplemental Figure 8B)**. Altogether, based on the expression of *Wt1* or *Pdgfra* and *Postn*, which broadly categorize cells into mesothelial- or mesenchymal-like states, we defined five major epicardial cell populations: Cluster 0: *Wt1^high^/Pdgfra^high^*; Cluster 1: *Wt1^-^/Postn^+^*; Cluster 2: *Wt1^high^/Msln^+^*; Cluster 3: *Wt1^low^/Pdgfra^high^*; and Cluster 4: *Wt1^mid^, Pdgfra^low^* **(Figure 2K)**. The remaining clusters were identified as part of the Wt1-lineage epicardial cell compartment but were classified as additional transcriptional states. These included a highly proliferative cluster (cluster 7) and lower-feature or transitional states (clusters 5, 6, and 8), rather than being categorized as the primary mesothelial or fibroblast-like populations **(Figure 2K)**. Collectively, these expression patterns support a model in which the epicardium undergoes transcriptional reprogramming towards a limited fibroblast phenotype following MI, while retaining a subset of epithelial-like cells.

### Differential gene expression of adult epicardial cell populations

To investigate significant cellular responses following myocardial injury, we conducted a differential gene expression (DEG) at two levels: globally across all epicardial-lineage cells by condition, and within individual epicardial cell clusters, comparing 7dMI versus Sham and 14dMI versus 7dMI. For both analyses, DEGs were set with a minimum expression threshold of 10% of cells and a log2 fold-change threshold of 0.25 for 7dMI versus Sham, and a minimum expression threshold of 5% of cells and a log2 fold-change of 0.10 for 14dMI versus 7dMI; genes with an adjusted p-value < 0.05 were considered significant. This two-level approach allowed us to capture broad injury-associated transcriptional shifts across the epicardial lineage while also resolving cluster-specific responses. In sham hearts, epicardial cells highly expressed adhesion- and barrier-associated genes such as *Bcam*, together with metabolic and transport genes (*Gpihbp1, Cd36, Aqp1*), and mesothelial identity gene markers (*Msln, Ly6a)* **(Figure 3A)**. High expression of antioxidant and redox-regulating genes such as *Gstm1* and *Selenop* further supports a quiescent, stress-protective epicardial state in the uninjured heart. In contrast, at 7dMI, epicardial cells upregulated genes involved in ECM remodeling and fibrosis (*Fn1*, *Plod2*, *Col5a1),* matricellular and angiogenic regulators (*Fstl1*, *Thbs1*, *Wisp1*), and EMT/cytoskeletal genes (*Ncam1*, *Tagln*, and *Csrp2)* **(Figure 3A)**. The expression of these genes is consistent with an epicardial transition from a homeostatic barrier layer toward a migratory, mesenchymal, matrix-producing, and pro-angiogenic reparative phenotype^4^. Analysis of DEGs in epicardial cells between 14dMI and 7dMI revealed a second phase of remodeling characterized by a global shift away from ECM/EMT-dominant signatures toward genes associated with immune modulation and injury resolution **(Figure 3B).** We observed high expression of transcripts linked to complement regulation and immune signaling (*C4b, Prelp, Clec3b, Nbl1, Sfrp2, Apoe, Pi16*), suggesting a functional transition from an early fibroblast-like reparative state at 7dMI to a more immune-interfacing epicardial state at 14dMI. Gene ontology (GO) analysis between conditions further illustrates this shift in function: terms upregulated in 7dMI vs Sham included Wnt signaling, EMT, angiogenesis, and fibroblast proliferation, whereas 14dMI vs 7dMI was enriched for humoral immune response, complement activation, IL-6 signaling, and leukocyte migration **(Figures 3C,3D)**.

**Figure 3.**
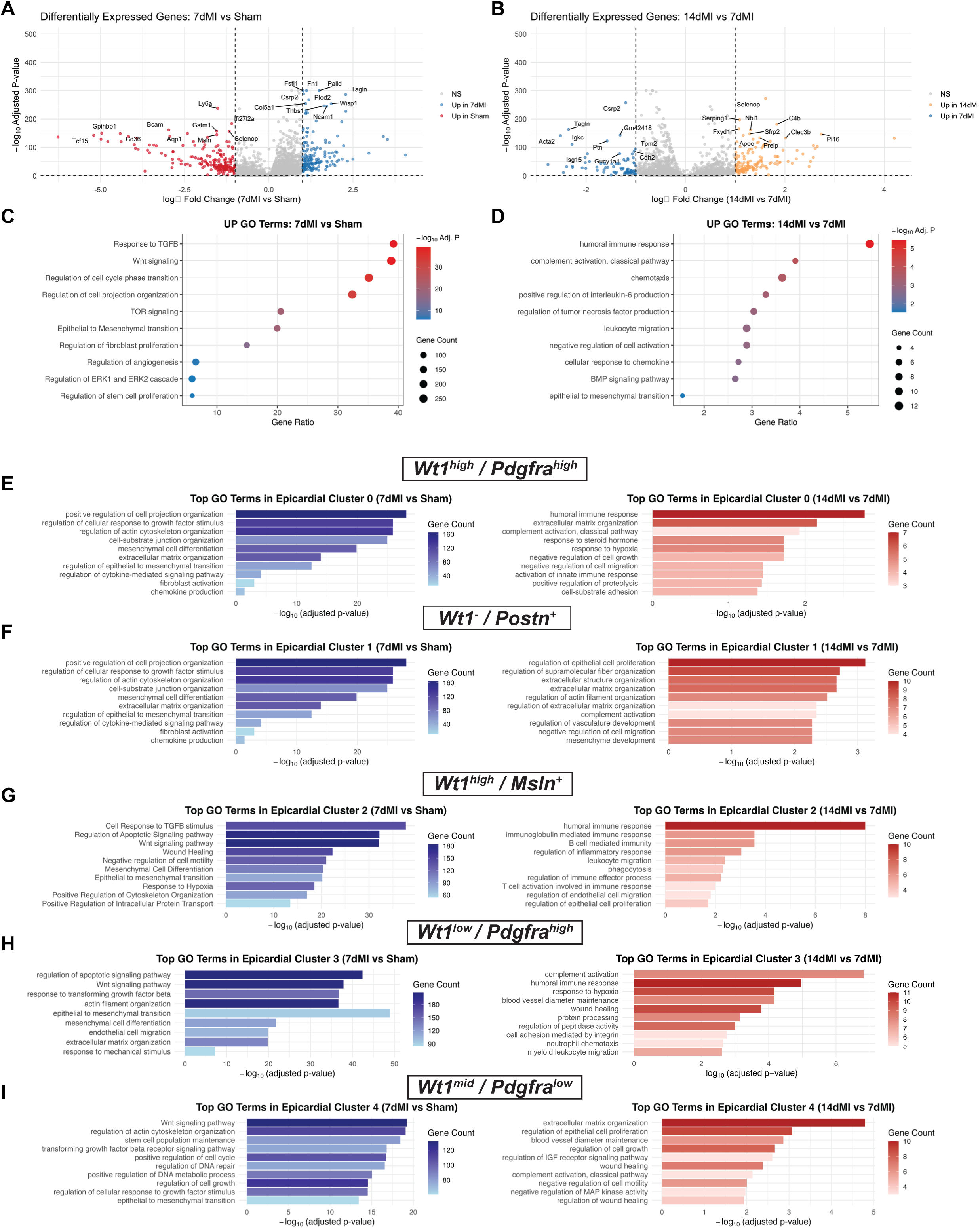
Acute myocardial ischemia triggers reparative gene programs in adult epicardial cells. **(A-B)** Volcano plots of differentially expressed genes (DEGs) comparing 7dMI vs Sham **(A)** and 14dMI vs 7dMI **(B)** acquired epicardial cells. Dashed lines indicate the statistical and fold-change thresholds used for identifying significant DEGs; the top 10 genes with the highest adjusted p-value are labeled. **(C-D)** Gene Ontology (GO) analysis of biological processes upregulated in 7dMI vs Sham **(C)** or 14dMI vs 7dMI **(D)** acquired epicardial cells. Dot size indicates the number of genes contributing to each term, and the dot color reflects enrichment significance (-log adjusted P value); the x-axis indicates gene ratio. **(E-I)** Cluster-specific GO analysis showing the top-enriched terms upregulated at 7dMI vs Sham (left, blue) and at 14dMI and 7dMI (right, red) in epicardial cells in Clusters 0-4. Bars represent -log(adjusted P value), with shading indicating gene counts contributing to each enriched term. Data are presented as biological replicates: N=5 for Sham, N=5 for 7dMI, and N=2 for 14dMI.

Cluster-specific GO term analysis highlighted the diverse biological roles of epicardial subpopulations during repair. Cluster 0 (*Wt1^high^/Pdgfra^mid^*) represents a population of fibroblast-like/transitioning epicardial-derived cells. GO terms upregulated at 7dMI vs. Sham included EMT, cell projection organization, and autophagy, suggesting an early activation of cell survival and matrix-remodeling programs **(Figure 3E).** By 14dMI, gene enrichment in Cluster 0 shifted toward humoral immune response and negative regulation of cell migration and growth, supporting a transition from structural remodeling to immunomodulatory and injury-resolution functions.

Cluster 1 (*Wt1^-^/Postn^+^)* is characterized by the enriched expression of *Postn*. GO terms enriched at 7dMI vs Sham included TGFβ signaling, ECM organization, actin filament organization, and fibroblast proliferation, all hallmarks of activated matrix-secreting fibroblasts **(Figure 3F)**. At 14 days post-MI, this subpopulation remained enriched for genes involved in ECM organization and remodeling, underscoring a persistent role for epicardial cells in engaging in fibrotic deposition and scar maturation during post-MI repair.

Cluster 2 (*Wt1^high^/Msln^+^*) displayed specific expression of epithelial-like identity genes, *Msln*^9^ and *Upk3b*^14^. GO terms enriched at 7dMI vs Sham included response to TGFβ stimulus, Wnt signaling, regulation of EMT, and wound healing **(Figure 3G)**. These pathways indicate that this epicardial subpopulation may primarily function as a signaling interface with the underlying myocardium and vasculature, supporting angiogenesis and tissue remodeling through paracrine cues. By 14dMI, Cluster 2 was strongly represented by immune-related GO terms, including T-cell activation, immune effector processes, and leukocyte proliferation, suggesting a late-stage role in coordinating immune cell recruitment and activation at the epicardial surface.

Cluster 3 (*Wt1^low^/Pdgfra^high^*) exhibits reduced *Wt1* expression but higher *Pdgfra*, representing a more differentiated mesenchymal subset compared to Cluster 0. At 7dMI, GO term enrichment included mechanical stress responses, ECM organization, and cytoskeletal remodeling, consistent with a mechanically engaged stromal compartment **(Figure 3H)**. By 14dMI, Cluster 3 transitioned toward complement activation, humoral immune response, and myeloid leukocyte migration, paralleling the immune-related shifts observed in other clusters and suggesting that differentiated epicardial-derived fibroblasts also contribute to shaping the late-stage immune-modulating/inflammatory milieu, as previously observed^7^.

Cluster 4 (*Wt1^mid^/Pdgfra^low^*) exhibited moderate *Wt1* expression with low *Pdgfra*. At 7dMI, GO terms included regulation of cell projection organization, ECM organization, and EMT, suggesting engagement in structural remodeling. By 14dMI, cluster 4 was enriched for wound healing, IGF signaling, and MAPK activity, implicating this subset as a stress-responsive, pro-tissue repair subpopulation.

GO term analysis of Clusters 5-8 further supports specialized roles for epicardial subpopulations. At 7dMI vs Sham, Clusters 5 and 6 were strongly enriched for genes related to TGF-β signaling, ECM organization, Wnt signaling, and wound healing **(Supplemental Figures 9A, 9B)**. Cluster 7 showed GO term enrichment for cell cycle, cytokinesis, and EMT, indicating a population of epicardial-derived cells dedicated to cell division **(Supplemental Figure 9C)**. Cluster 8 shared many terms with Clusters 5 and 6, including ECM organization and Wnt signaling, but also displayed GO term enrichment related to the regulation of protein secretion, suggestive of a secretory, matrix-producing phenotype **(Supplemental Figure 9D)**.

In all epicardial subpopulations, the early responses observed after 7dMI were mainly influenced by developmental pathways, EMT processes, and matrix remodeling pathways, as evidenced by increased upregulated DEGs in 7dMI clusters compared to sham **(Supplemental Figure 9E).** In contrast, the late responses noted at 14dMI were characterized by immune regulatory programs and a large downregulation of 7dMI gene programs **(Supplemental Figure 9E)**. Together, these observations suggest that epicardial cells undergo a temporary activation resembling a developmental state following ischemia. This temporal and cluster-specific reprogramming mirrors the canonical sequence of post-injury repair, highlighting how epicardial cells adopt distinct but complementary functional states to orchestrate cardiac remodeling and resolution following MI.

### Cell-cell communication between epicardial cell subpopulations

To further define how MI alters epicardial paracrine signaling networks, we performed ligand-receptor-based CellChat analysis^28^. We found that the mesothelial-like (*Wt1^high^/Msln^+^)* and fibroblast-like (*Wt1^high^*/*Pdgfra^mid^* and *Wt1^-^/Postn^+^*) epicardial states function as major communication hubs, exhibiting high incoming and outgoing interaction strengths, whereas clusters 5, 6, and 8 showed comparatively weak cell-cell signaling **(Figures 4A, 4B, and Supplemental Figure 10)**. Pattern recognition of pathway-level signaling further indicated that *Wt1^high^/Msln^+^*, *Wt1^-^/Postn^+^,* and *Wt1^high^/Pdgfra^mid^* clusters are a dominant source of ECM and growth factor family of signaling, related to TGFβ, SEMA3, TENASCIN, IGFBP, NOTCH, CXCL, FLRT, NETRIN, WNT, and non-canonical WNT, and VEGF **(Figure 4C and Supplemental Figures 11A-F)**. Wnt signaling has been implicated in injury-induced epicardial activation, in which β-catenin-dependent signaling promotes epicardial expansion and EMT after ischemic cardiac injury^12^. In our dataset, both canonical and non-canonical WNT pathway activity were highly enriched in the mesothelial-like *Wt1^high^/Msln^+^* cells **(Supplemental Figure 11B,C)**, suggesting that Wnt signaling regulates the cell-state transitions observed in our post-MI epicardium. Altogether, the mesothelial-like *Wt1^high^/Msln^+^* cluster exhibited a high degree of incoming signals from several reparative-like signaling inputs, including VEGF **(Supplemental Figure 11D)**, suggesting a highly receptive epicardial state **(Figure 4C)**.

**Figure 4.**
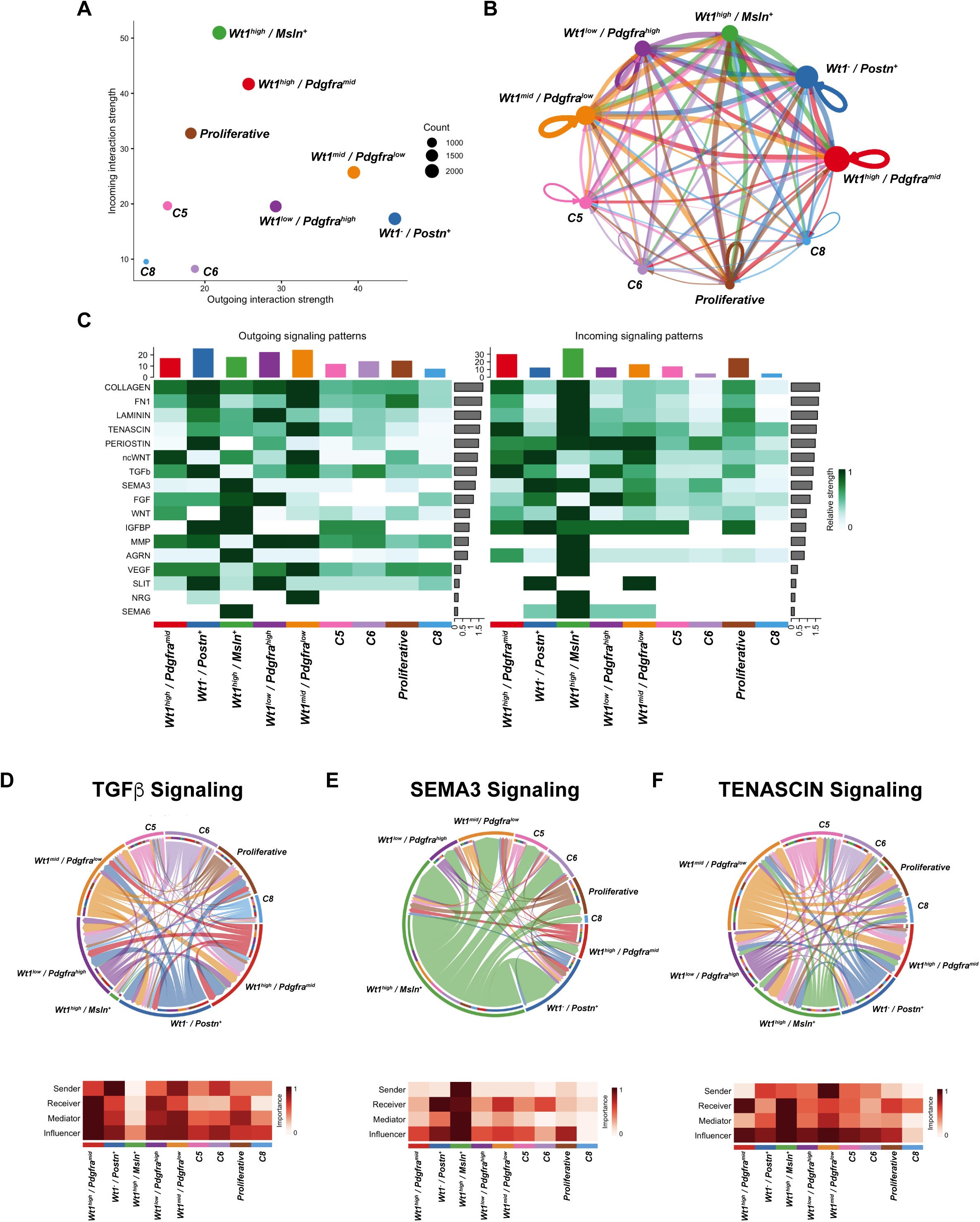
A dynamic network of communication between epicardial cells. **(A)** Scatter plot summarizing global communication roles for each epicardial cell cluster, plotting outgoing (sender) interaction strength versus incoming (receiver) interaction strength inferred by CellChat. Point size reflects the number of cells in each cluster. **(B)** Circular network visualization of the aggregate ligand-receptor communication network among epicardial clusters. Directed edges represent inferred intercellular interactions; edge thickness reflects interaction strength, and node size corresponds to cluster size. **(C)** Heatmaps of pathway-level activity, showing the relative contribution of indicated signaling pathways to outgoing (left) and incoming (right) communication across epicardial clusters (color scale indicates relative signaling strength). **(D-F)** Representative pathway-focused networks for TGFβ **(D)**, SEMA3 **(E)**, and TENASCIN **(F)** signaling. The top panels show chord diagrams depicting the direction and magnitude of communication between sender and receiver states within each pathway. Bottom: Signaling role heatmap summaries (sender, receiver, mediator, influencer) for each epicardial state within the corresponding pathway. Data are presented as biological replicates: N=5 for Sham, N=5 for 7dMI, and N=2 for 14dMI.

After myocardial injury, TGFβ ligands are induced, and canonical SMAD signaling is enhanced within the injured heart, highlighting the central role of TGF-β in facilitating tissue repair^13^. *Wt1^high^/Pdgfra^mid^* and *Wt1^-^/Postn^+^*populations represent the principal senders and influencers, expressing high levels of TGFβ ligands that impact downstream signaling in other epicardial populations, including *Wt1^low^/Pdgfra^high^* and the proliferative subset **(Figure 4D)**. In contrast, the mesothelial-like *Wt1^high^/Msln^+^* subpopulation displayed little to no expression of TGFꞵ signaling components and exhibited very minimal cross-talk with other epicardial cell populations **(Figure 4D)**. Notably, SEMA3 signaling was increased in *Wt1^high^/Msln^+^* mesothelial-like cells, which acted as the primary communication hub and key senders/mediators to mesenchymal and proliferative populations **(Figure 4E).**

Tenasin-C (TNC) is induced after MI and becomes spatially restricted to the infarct border zone^29^; therefore, we examined TNC signaling as a candidate ECM cue coordinating injury-induced epicardial state transitions, including EMT-like remodeling. TNC signaling was most prominently produced by *Wt1^-^/Postn^+^*and *Wt1^mid^/Pdgfra^low^* fibroblast-like cells and was widely distributed across all epicardial states **(Figure 4F)**, consistent with prior work implicating TNC as a pro-migratory and ECM-regulatory cue^30^. Altogether, these data indicate that post-MI remodeling is coordinated by both mesothelial- and fibroblast-like epicardial cells that express differential communicative factors, such as TGFꞵ, semaphorins, and tenascin.

### Expression of TGFb, WNT, EMT-associated, and Semaphorin signaling genes in epicardial cells during ischemic progression

As our CellChat analysis identified TGFꞵ, WNT, and Semaphorin as major communication axes between epicardial subpopulations after MI, we further examined individual genes in these pathways and their downstream effectors that regulate epicardial cell-specific function and/or activation during cardiac repair. Our data revealed robust induction of pathways in adult epicardial cells after MI, including TGFβ ligands, receptors, and SMAD pathway components, which were broadly upregulated at 7dMI across fibroblast-like and proliferative clusters, with *Tgfbr1* and Tgfb2 showing higher expression in Clusters 0 (*Wt1^high^/Pdgfra^high^*), 3 (*Wt1^low^/Pdgfra^high^*), and 4 (*Wt1^mid^, Pdgfra^low^*), respectively **(Figure 5A-C)**.

**Figure 5.**
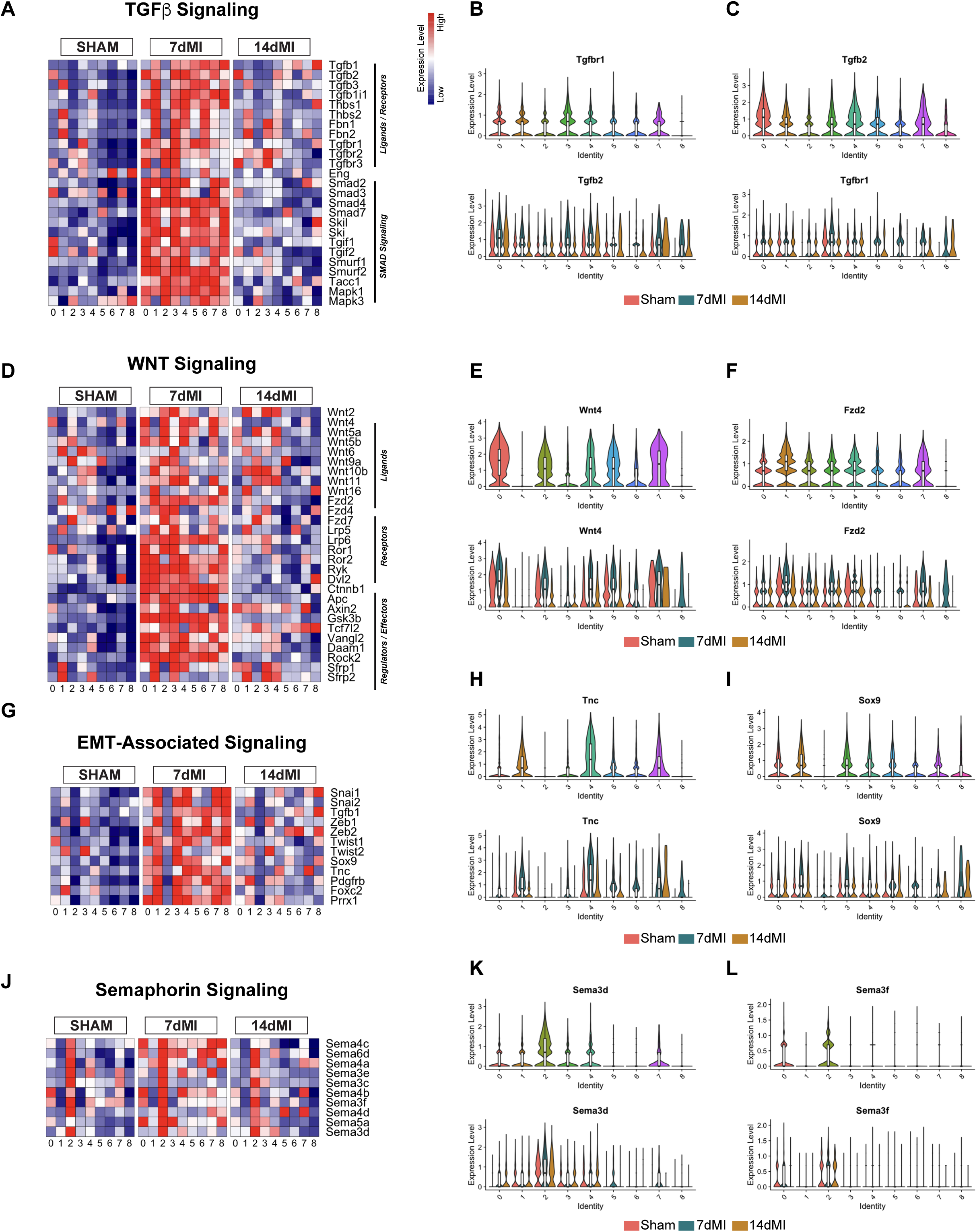
Ischemia-driven alterations in gene expression highlight reparative signaling programs. Heatmap of ligand-receptor interaction potential scores between epicardial ligands and cognate receptors across conditions belonging to **(A)** TGFβ signaling. Violin plots representing **(B)** *Tgfbr1* and **(C)** *Tgfb2* expression amongst cell clusters and conditions. Expression heatmap of **(D)** WNT signaling, **(E)** *Wnt4*, and **(F)** *Fzd2*. Expression heatmap of **(G)** EMT-associated signaling, **(H)** *Tnc*, and **(I)** *Sox9*. Expression heatmap of **(J)** Semaphorin signaling, **(K)** *Sema3d*, and **(L)** *Sema3f*. Data are presented as biological replicates: N=5 for Sham, N=5 for 7dMI, and N=2 for 14dMI.

As Wnt signaling is a critical regulator of epicardial development and adult tissue repair^31^, we interrogated the expression of Wnt ligands, receptors, and downstream effectors across conditions. At 7dMI, non-canonical ligands (*Wnt4, Wnt5a, Wnt11*) and receptors such as *Fzd2* were robustly induced with preferential enrichment in cluster 0 (*Wt1*^-^/*Postn^+^* cells) and the ligands being expressed in *Wt1*-expressing epicardial cells **(Figure 5D-F)**. By 14dMI, expression of these genes was generally diminished, indicating a defined early post-infarct window of Wnt pathway activation **(Figure 5D)**. Together, these data support a role for Wnt and TGFβ signaling in coordinating epicardial activation and paracrine communication during early cardiac repair. EMT-associated transcription factors and effectors (*Snai1/2, Zeb1/2, Twist1/2, Sox9, Tnc, Pdgfrb, Prrx1*) were expressed at relatively lower levels in sham epicardial cells compared to cells from MI conditions, and were noticeably upregulated at 7dMI across multiple states, as evidenced by *Tnc* enrichment in clusters 1 *Wt1^-^/Postn^+^* and 4 (*Wt1^mid^/Pdgfra^low^*) and *Sox9* being broadly upregulated across cell clusters during this injury phase, except in mesothelial cluster 2, suggesting that most adult epicardial cells upregulate EMT genes during acute ischemic injury. **(Figure 5G-I)**. In some cases, EMT-associated genes persisted at 14dMI, suggesting that subsets of epicardial cells maintain a mesenchymal-like state during late-stage post-MI cardiac repair **(Figure 5G)**.

Notably, we observed minimal expression of EMT-associated genes in the mesothelial-like *Wt1^high^/Msln^+^* population, suggesting that this subpopulation of epicardial cells undergoes minimal conversion to a mesenchymal/fibroblast-like phenotype. Indeed, the mesothelial-like *Wt1^high^/Msln^+^* population showed high expression of semaphorin genes, such as *Sema3d* and *Sema3f,* at 7dMI **(Figure 5J-L)**, consistent with CellChat analyses, suggesting that guidance cues are activated in response to injury to modulate remodeling and intercellular signaling in surface epicardial cells.

Although studies of semaphorin function in adult injury contexts remain sparse, investigations during development have shown that *Sema3c* is required for proper cardiac outflow tract septation and vessel development^32^, while *Sema3f* is essential for heart chamber formation^33^.

At 14 days post-injury, the expression of many genes associated with injury was reduced, indicating a shift from acute activation to a more stable remodeling phase. Consistent with this, the expression of a broad array of ECM genes, including collagens (*Col1a1, Col1a2, Col3a1*), matricellular proteins (*Fn1, Postn, Sparc, Thbs1*), and ECM remodelers (*Mmp2, Mmp9, Adamts* family) was strongly induced at 7dMI and partially resolved by 14dMI **(Supplemental Figure 12A)**. Cluster-specific expression of Fibroblast activating protein *(Fap)* marked *Wt1*^-^/*Postn^+^*myofibroblast-like cells, while *Fn1 and Thbs1* were more broadly distributed and sustained **(Supplemental Figures 12A-D)**. In contrast, immune response genes, including complement components (*C4b, C1qa, C1qc)* and regulators such as *Svep1* and *Adm*, were highly expressed at 14 days post-MI **(Supplemental Figures 12E-H)**, suggesting a shift in epicardial cells from involvement in matrix remodeling to immune modulation. In addition to these gene programs, epicardial clusters also exhibited selective expression of reported tissue repair mediators, including *Fstl1* and *Tmsb4x*, which have been shown to regulate cardiac regeneration and vascular growth through *Vegfa, Slit2,* and *Cxcl12,* as well as EMT and fibrotic processes through *Ccm2* and *Col5a1* **(Supplemental Figures 12A-D)**. Altogether, these results demonstrate that epicardial cells coordinate multiple signaling pathways, including semaphorins, EMT regulators, tissue repair, chemokines, inflammation, and Wnt pathways, in a stage-specific manner to orchestrate myocardial tissue remodeling and repair following injury.

### Investigating the distribution of epicardial cells during heart remodeling after myocardial infarction

To observe the localization of epicardial cells in response to MI, we performed immunohistochemistry and fluorescence in situ hybridization (FISH) on *Wt1^CreERT2/+^; R26^tdTomato/+^* hearts subjected to sham or MI, following tamoxifen administration as previously detailed **(Figure 1B)**. First, we observed tdTomato expression on the epicardial surface of sham hearts, with a pronounced epicardial expansion in MI conditions **(Figure 6A-C)**. Notably, we found that several tdTomato^+^ epicardial cells in sham conditions co-expressed PDGFRα-GFP reporter, which increased at 7dMI and decreased at 14dMI **(Figure 6A-C)**. Utilizing FISH, *Msln* was confined to the outermost mesothelial sheet of the uninjured heart **(Figure 6D)**, demarcating the epicardial surface, consistent with previous studies^9^. Additionally, in the sham heart, *Wt1-*lineage (*tdTomato*) cells form a thin continuous layer along the outer surface, with a sparse number of cells in the underlying myocardium that do not express *Postn* **(Figures 6D)**. At 7dMI, the *tdTomato*-expressing epicardial region dramatically thickens and organizes into a laminar arrangement: a reduced *Msln^+^* surface layer, an overlapping as well as subadjacent *tdTomato*^+^ band, and a broad *Postn^+^* compartment extending into the subepicardium and infarct border zone **(Figure 6E)**. Additionally, the *Postn^+^* cells are robustly induced, and densely populate the subepicardial infarct border zone, whereas *tdTomato^+^* cells continue to line the surface and show limited co-expression of *Postn,* suggesting a partial contribution of *Wt1*-lineage cells to the activated fibroblast pool **(Figure 6E)**. At 14dMI, we continue to observe an expanded *tdTomato^+^* sheet along the surface and infarct edge, and the *Postn^+^* compartment extends deeper into the fibrotic areas **(Figure 6F)**. By 28dMI, signals associated with epicardial activation partially resolve: *Postn* intensity diminishes; *tdTomato*^+^ cells are located on the surface of the heart, many *Wt1*-lineage cells persist in the subepicardium, and the expression of *Msln* returns to levels similar to those observed in sham hearts, suggesting that some cells may revert to a mesothelial-like state **(Figure 6G)**. Taken together, these data show that MI triggers a dynamic expansion of the epicardial region, in which *Wt1-*lineage cells persist as a superficial layer with partial overlap with an underlying PDGFRα*^+^ and Postn^+^* cellular compartment that peaks at 7-14 days and reverts to homeostatic levels by 28 days, consistent with a proposed time course of early epicardial activation and partial EMT, followed by remodeling and resolution.

**Figure 6.**
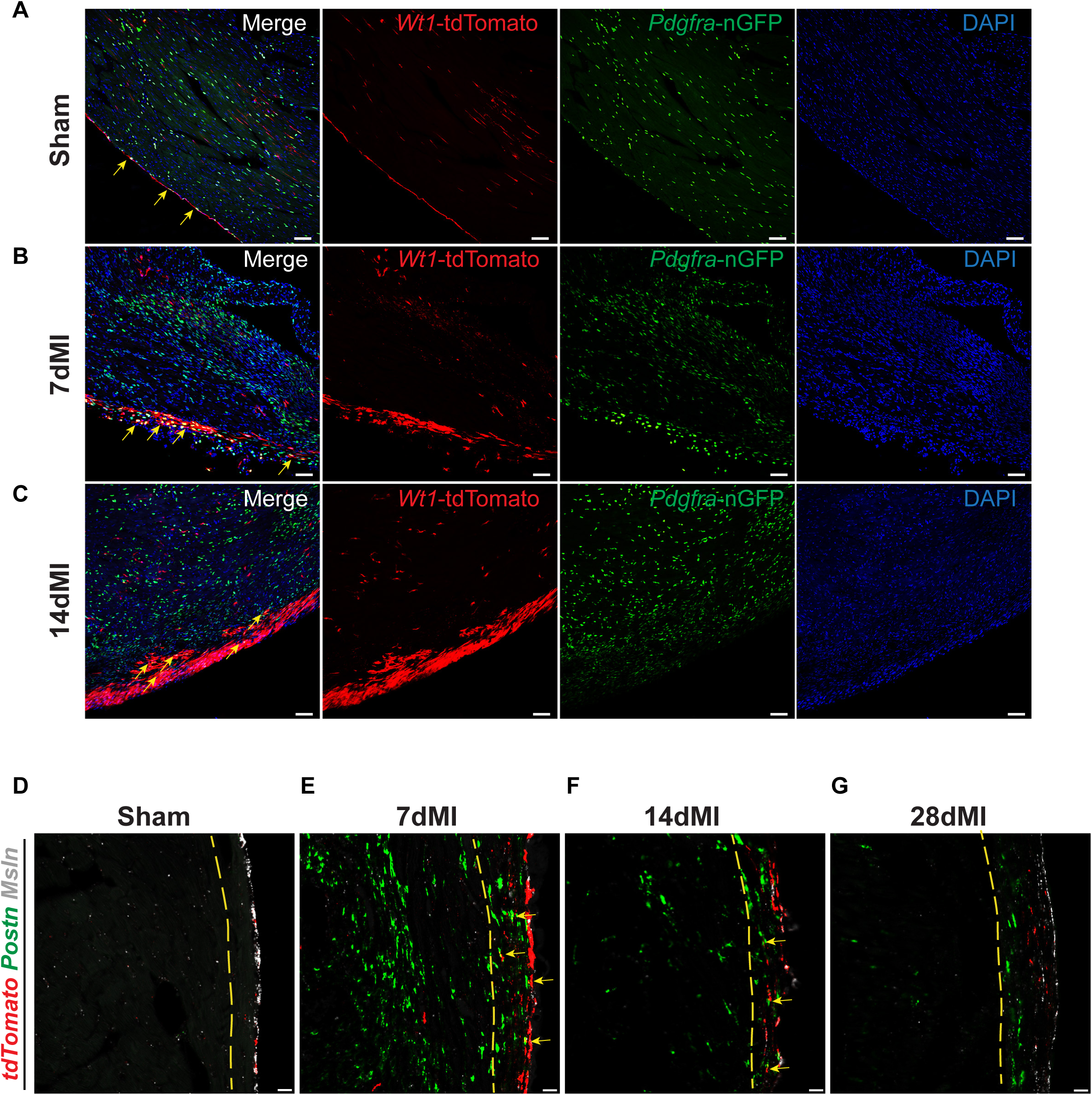
Ischemia leads to a transient expression of fibroblast markers in epicardial cells. **(A-C)** Representative immunofluorescence images of *Wt1^CreERT2/+^; R26^tdTomato/+^; Pdgfra^nGFP/+^* subjected to sham and 7 or 14 days of MI. Arrows highlight co-expression of tdTomato (red) and GFP (green). Blue = DAPI nuclei. Scale bar: 50μm. **(D-G)** Fluorescence in situ hybridization images of left ventricular sections from sham, 7dMI, 14dMI, and 28dMI showing the spatial distribution of lineage-traced epicardial cells (*TdTomato*, red) amongst injury-induced Periostin (*Postn*) expression. Mesothelin (*Msln*) was used to visualize the intact epicardium. The yellow dashed line indicates the subepicardial boundary; Arrowheads indicate *Wt1*-tdTomato^+^ epicardial cells co-localizing with *Postn*. Blue = DAPI nuclei. Scale bar: 50μm. Images are from a single biological animal, and all staining was performed on at least N=3 biological replicates.

### Adult epicardial cells increase the expression of EMT-related genes after injury beyond the levels observed in fetal epicardial cells

To determine how adult uninjured (sham) and injured (7dMI, 14dMI) epicardial cells differ from epicardial cells during cardiac development, we integrated our adult epicardial cell scRNA-seq data with transcriptomics previously conducted in *Wt1*-lineage epicardial cells from key developmental timepoints at embryonic day (E)12.5 and E16.5 **(Supplemental Figures 13A-C)**^34^. Each sample was processed as an individual Seurat object and normalized independently using SCTransform. To enable comparison across datasets, all samples from both studies were then jointly integrated using Seurat’s SCT-based integration workflow, including shared feature selection, anchor identification, and integration into a common low-dimensional space. Following integration, we observed *Wt1* expression across the integrated epciardal cell atlas **(Figures 7A, 7B)**. Pseudobulk PCA separated fetal from adult epicardial profiles along the dominant axis of variation (PC1), displaying that adult sham and post-MI samples remained transcriptionally similar as compared to fetal epicardial cells **(Figure 7C)**. Additionally, Pearson correlation analysis further supported this distinction: adult pseudobulk transcriptional profiles were highly correlated across sham, 7dMI, and 14dMI, whereas fetal timepoints E12.5 and E16.5 were transcriptionally correlated with each other **(Figure 7D)**. Gene set comparisons reveal that 7dMI promotes a mesenchymal activation/EMT-like program, surpassing fetal levels, with relatively higher expression of key EMT-associated regulators and effectors (*Snai1/2, Zeb1/2, Twist1/2, Sox9, Tnc*) compared to both sham and fetal epicardium, which then subsides by 14dMI **(Figure 7E)**. In contrast, embryonic-derived epicardial cells, particularly at E12.5, showed higher expression of canonical epicardial gene markers, including *Wt1*, *Msln*, *Upk3b*, *Tbx18*, and *Aldh1a2* **(Figure 7E)**. Despite the developmental age of epicardial cells, fetal and adult cells expressed the canonical developmental/epithelial epicardial marker *Wt1* **(Figure 7B,7F)**, as well as quiescent fibroblast marker *Pdgfra* **(Figure 7J)**. However, other epicardial markers, including *Msln*, *Upk3b*, and *Tbx18*, were more highly expressed in epicardial cells from the adult sham heart and from the E12.5 and E16.5 developmental stages, and were downregulated following MI **(Figures 7G-I)**. Adult epicardial cells exhibited higher levels of *Postn* than embryonic-derived epicardial cells, and notably, *Postn* was transiently induced in epicardial cells during acute MI **(Figure 7K)**. The data suggest that MI triggers EMT, which may serve as an adaptive response to cardiac injury. Furthermore, the timing of EMT across developmental stages is highly tunable and closely corresponds to the differentiation of epicardial cell populations in the developing heart. This suggests potential opportunities for targeted therapeutic strategies to improve cardiac repair after MI, underscoring the need for further investigation into EMT dynamics in adult cardiac recovery.

**Figure 7.**
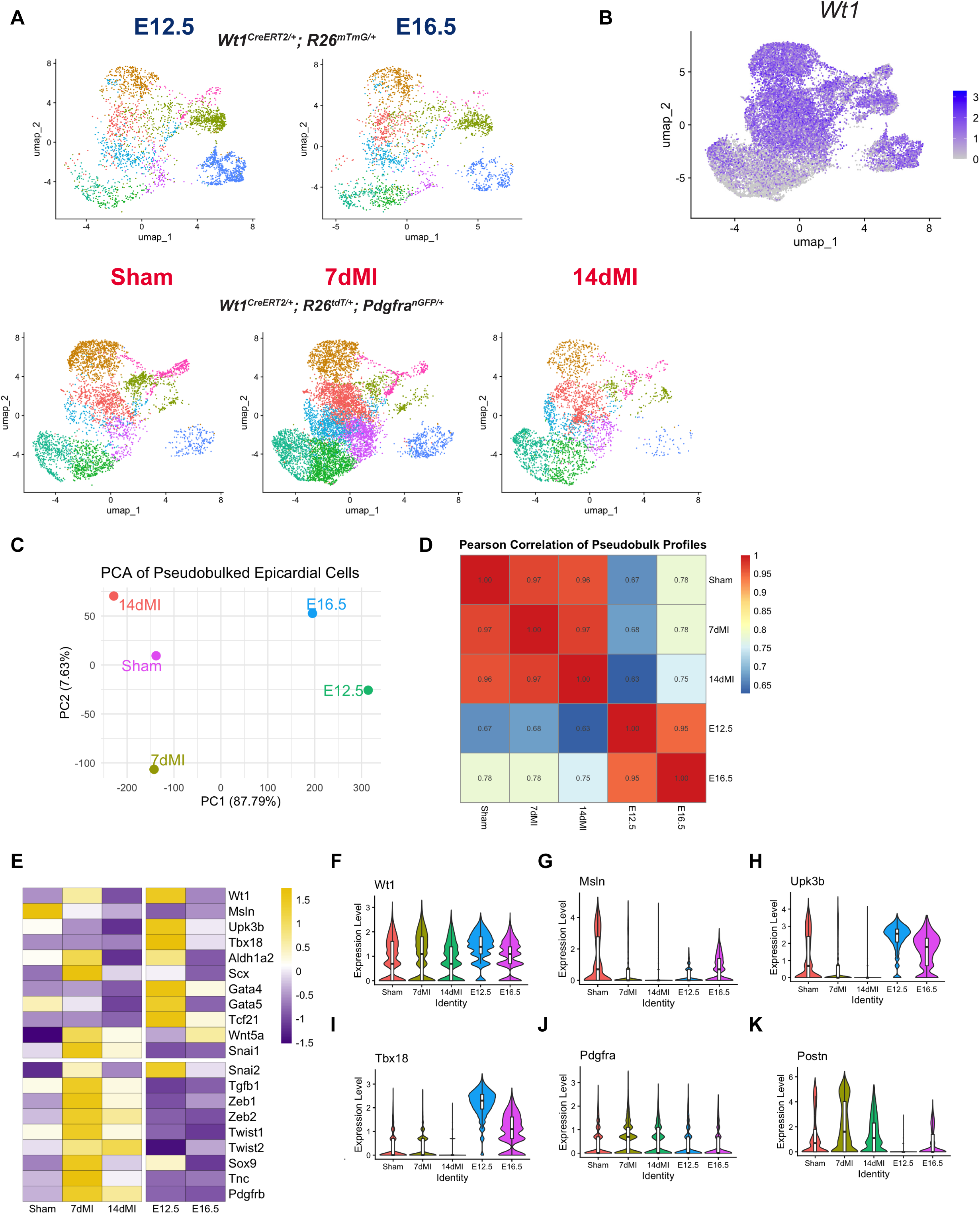
A comparison of epicardial dynamics during cardiac development and acute ischemia. **(A)** UMAP of lineage-traced epicardial cells acquired from fetal hearts at embryonic day (E) 12.5 and E16.5 from *Wt1^CreERT2/+^; R26^mTmG/+^* mice and adult hearts following Sham, 7dMI, or 14dMI from *Wt1^CreERT2/+^* ; R26^tdTomato/+^; Pdgfra^nGFP/+^ mice. **(B)** FeaturePlot showing *Wt1* across the integrated fetal-adult epicardial cell UMAP. **(C)** Principal component analysis (PCA) of pseudobulked epicardial transcriptomes for each condition/time point. **(D)** Heatmap of Pearson correlation coefficients computed between pseudobulk expression profiles across all conditions, quantifying similarity between fetal and adult epicardial programs. **(E)** Scaled expression heatmap of selected epicardial/mesothelial identity and developmental/EMT-associated genes across conditions. **(F-K)** Violin plots of epicardial/mesothelial and fibroblast markers across fetal and adult heart conditions. Data are presented as biological replicates: N=5 for Sham, N=5 for 7dMI, and N=2 for 14dMI.

## DISCUSSION

Here, we present a single-cell framework for understanding how the adult epicardium responds to both acute and chronic stages of ischemic injury. Previous studies have demonstrated that acute MI (5 days post-MI) triggers epicardial proliferation and limited subepicardial migration, contributing to the formation of a thickened surface layer that primarily engages in post-reparative paracrine signaling^4^. However, the profiling of adult epicardial cells during the later stages of cardiac injury has not yet been clearly defined. To build on our molecular understanding, we examined the time course of adult epicardial cell reactivation by performing single-cell RNA sequencing of *Wt1*-lineage adult epicardial cells isolated from 7-day- and 14-day-injured hearts and compared them with non-injured controls. Condition-level differential gene expression and pathway enrichment analyses revealed that 7 days post-MI, *Wt1*-lineage epicardial cells displayed enrichment for EMT and ECM-remodeling programs, along with pro-reparative pathways associated with angiogenesis, cardiomyocyte growth, and fibrotic remodeling. By 14 days post-MI, the epicardial cell response to injury shifted toward immune- and complement-regulatory programs, with reduced expression of early reparative gene signatures. Analysis of epicardial cell heterogeneity further revealed a persistent epithelial/mesothelial population, transitory and mature mesenchymal cell subtypes, and a minor cluster of proliferative epicardial cells. Overall, this study provides a platform for analyzing the temporal and cellular heterogeneity in the epicardium and its response to ischemia, offering insights into the likely diverse range of reparative and immune-modulating functions of epicardial cells.

### Heterogeneity and functional division within the epicardial compartment

In the current study, we resolved 9 adult epicardial cell clusters derived from the *Wt1* Lineage, focusing on the following major epicardial states: epithelial/mesothelial-like cells (*Wt1^high^/Msln^+^*), mesenchymal cells (*Pdgfra^low-high^*), and transitional/proliferative cellular intermediates. These broad cell states are not entirely de novo, but instead correspond to conserved or recurrent epicardial programs described during development and adult cardiac injury. Prior work has shown that MI induces the re-expression of embryonic epicardial cell markers, mesenchymal/paracrine programs, while single-cell analyses have further demonstrated substantial heterogeneity within the post-MI epicardial stromal compartment. Our study confirms these major adult epicardial cell states, while extending previous work by defining their *Wt1-*lineage composition and temporal remodeling across sham, 7dMI, and 14dMI hearts. At 7dMI, mesenchymal-like clusters (Clusters 1/3/4) upregulate canonical matrix and matricellular genes (*Fn1*/*Thbs1*/*Col5a1)*, cytoskeletal genes, and EMT transcription factors (*Snai1/2, Zeb1/2*), consistent with increased migratory capacity and matrix deposition. This data is consistent with observations in adult zebrafish regeneration, where activated epicardial cells secrete paracrine factors that induce resident cardiomyocytes to re-enter the cell cycle^35^. Additionally, zebrafish *tcf21^+^* epicardial cells migrate to the site of injury, re-express developmental programs, undergo EMT, and differentiate into cells of the non-CM lineage^10,22^. In contrast, in mammals, multiple fate-mapping studies indicate that after MI, *Wt1*-lineage epicardial-derived cells accumulate on the heart surface, exhibit limited capacity to differentiate into mesenchymal derivatives (fibroblasts/smooth muscle cells), and exert pro-reparative effects through paracrine signaling, despite limited myocardial invasion^36^. Consistent with this observation, FISH identified a superficial *Wt1*-lineage band that is preserved through 28dMI, which is subjacent to the *Pdgfra*^+^/*Postn*^+^ compartment that emerges between 7 and 14dMI and regresses by 28dMI, with little to no adult epicardial cells present in the intramyocardial regions. This suggests that MI induces transient EMT in the murine heart, which does not promote significant migration and/or differentiation of adult epicardial cells, but rather enhances their paracrine signaling.

By contrast, the epithelial-like cluster 2 retains epithelial gene signatures (*Msln, Upk3b),* displays limited EMT-associated activation, yet is enriched for Wnt signaling and semaphorin-associated genes, consistent with a paracrine/coordinating role that does not require full mesenchymal conversion. Although not studied in the adult heart, semaphorin signaling may provide an epicardial “interface” function, given the established roles of epicardial/pericardial-derived semaphorins in cardiovascular patterning and vessel/lymphatic guidance^32,33,37^. Semaphorin signaling showed higher expression in the *Wt1^high^/Msln^+^,* suggesting that guidance cues may contribute to epicardial cell positioning, boundary maintenance, and intercellular communication during early remodeling. This interpretation is supported by genetic and functional studies showing that Semaphorin3E-PlexinD1 signaling is required for ventricular compaction, and inhibition of Sema3E-PlexinD1 after MI enhances lymphangiogenesis and improves cardiac function after MI^37^. These data suggest compartmentalized functions in the post-infarct epicardium: a surface epithelial/mesothelial subset specialized for intercellular signaling and environmental sensing, and a mesenchymal subset specialized for transient ECM remodeling and structural repair.

Our observed epicardial cell profiles during late cardiac repair (14dMI) are consistent with the concept that the adult epicardium functions as a reactive signaling barrier that adapts to a leukocyte-rich environment during ischemia, rather than serving as a major source of immune cells^7,9,20^. Indeed, prior work has demonstrated that the adult epicardium contains resident CD45^+^ hematopoietic cells and that injury-reactivated Wt1^+^ epicardial cells do not acquire the expression of immune cell markers within the reactivated epicardium^38^. In line with this, we did not detect *Ptprc* (CD45) expression within our *Wt1*-lineage epicardial clusters, supporting a model in which immune cells are recruited to and signaling within an expanded supracardial niche, but are not generated by the transdifferentiation of *Wt1*-lineage epicardial cells after injury. Notably, Complement C3 (C3) has been identified as a marker of the mature epicardium, in addition to the expression of genes associated with ECM^20^. Additionally, the aged adult epicardium exhibited increased signaling related to immune regulation compared to fetal epicardial cells^14^, supporting the notion that epicardial cells following 14 days of ischemia may revert to a more mature/aged state during chronic stages of ischemic disease progression.

### Epicardial signaling pathways that coordinate remodeling and repair

Cell-cell communication analysis indicates that the post-MI epicardium serves as both a source and a target of conserved developmental signaling pathways involving TGFβ, canonical/non-canonical Wnt, axon-guidance cues (semaphorins), and matricellular ECM regulators (TNC), which coordinate epithelial and epicardial-based activation, EMT, and remodeling^39^. TGFβ signaling is a central driver of EMT during development and a critical player induced in the infarcted heart to initiate remodeling^13,40^. In embryogenesis, ALK5 (TGFβ type 1 receptor)-dependent signaling is required for epicardial EMT and expansion, and disruption of epicardial ALK5 results in coronary/myocardial growth defects^41^. Indeed, adult epicardial cells retain strong TGFβ responsiveness in vitro, as treatment of human-derived cells with TGFβ3 induced EMT-like transformation, whereas ALK5 inhibition prevented EMT^42^. However, the extent to which epicardial cells serve as an endogenous source of TGFβ ligands or respond to TGFβ produced by fibroblasts, vascular cells, or infiltrating immune cells remains unclear, as we primarily focused on epicardial cell transcriptomics. Our signaling results are most consistent with a framework in which epicardial cells are TGFβ-competent responders that also participate in the expression/secretion of TGFβ-associated signaling ligands with the MI-induced epicardium.

Downstream of TGFβ signaling, EMT-associated transcriptional regulators (Snai1/2, Zeb1/2, Twist1/2, Sox9) were found to be highly expressed by all epicardial cell clusters, with the lowest relative expression in the mesothelial/epithelial-like *Wt1^high^/Msln^+^* population. Notably, recent work has demonstrated that Sox9 in the epicardium promotes invasive behavior and a fibroblast-like phenotype, supporting the idea that Sox9^+^ epicardial states represent a pro-migratory phenotype^43^. Prior reviews have highlighted the association of TNC with cell migration and EMT, with TNC being expressed in the proepicardial organ near migrating cells, a critical site of EMT transition^44,45^. Other evidence shows that TNC is induced during injury and tissue remodeling, demarcating border zones and facilitating the recruitment of myofibroblasts^46,47^. We also observed induction of tenascin-linked matricellular programs, with mesenchymal/fibroblast-like epicardial cells expressing TNC and TNXB, suggesting that ECM composition may be an injury-induced input that feeds back onto epicardial identity and EMT competence. In our dataset, the expression of TGFβ and Wnt-associated modules, along with the induction of EMT transcription factors and ECM components, is consistent with the selective reactivation of developmental epicardial signaling programs during adult cardiac injury. However, comparison with E12.5/E16.5 epicardial cells indicates the adult post-MI response does not represent a uniform return to an embryonic epicardial state. Rather, adult MI induces an injury-adapted transcriptional program in which developmental EMT/ECM modules are reployed in the context of ischemic remodeling, subepicardial expansion, and scar maturation. This interpretation is further supported by our fetal/adult epicardial cell integrated analysis, which showed that adult MI reactivates selected developmental markers and EMT-associated programs but remains transcriptionally distinct from the E12.5/E16.5 epicardium.

Consistent with the general phases of the post-MI injury response^21^, immune and immune-modulatory programs dominated GO enrichment of upregulated DEGs in 14dMI versus 7dMI epicardial cells. After MI/reperfusion, neutrophils have been observed not only within the infarct but also on the epicardial surface^35,48^, suggesting the epicardium functions as an immune cell reservoir. Additionally, epicardial Hippo pathway effectors YAP/TAZ have been shown to promote an immunosuppressive program after MI by recruiting regulatory T cells^7^. Our observation that immune-modulatory programs, such as complement genes/chemotactic factors, become enriched at 14dMI is consistent with accumulating evidence that the epicardium shapes or responds to post-infarct inflammatory resolution and a leukocyte-rich subepicardial niche. Altogether, these data support a two-phase model in which early epicardial activation (7dMI) is enriched for EMT/ECM/guidance modules that organize the reparative niche, whereas later epicardial states (14dMI) engage in immune-regulatory signaling to shape resolution and scar maturation.

### Comparison of epicardial cell function in fetal and adult hearts

Our integrated fetal (E12.5/E16.5) versus adult (sham/MI) analysis refines the prevailing view that the adult post-MI epicardium “recapitulates development.” While adult injury reactivates a subset of developmental epicardial markers (*Wt1, Tbx18, Aldh1a2)*^4^, this injury response is not a uniform return to a fetal epicardial identity/state. Instead, MI induces a stress-adapted activation program characterized by robust EMT/mesenchymal activation, which, in aggregate, can exceed fetal epicardial levels for core EMT regulators/effectors. This distinction is consistent with studies showing that the fetal epicardium harbors migratory/fibroblast-like populations and pro-angiogenic paracrine programs, whereas the adult epicardium is comparatively mesothelial, immune-responsive, and less overtly regenerative^14^. Our data suggest that adult MI elicits a stress-adapted EMT-like state, potentially tuned for rapid cardiac sealing, subepicardial expansion, scar remodeling, and immunomodulation, rather than a developmental epicardial state optimized for growth and vascular morphogenesis.

In conclusion, our investigation of the adult murine epicardium under ischemic stress highlights the limited regenerative capabilities of epicardial cells. Unlike complete cardiac repair models in other organisms, which exhibit similar developmental and reparative processes, the mammalian epicardium is not governed by conserved gene regulatory mechanisms^49^. This study presents a temporal analysis of adult Wt1-lineage epicardial cells during cardiac injury and repair, highlighting previously unknown diversity and functional specialization. Our findings enhance the understanding of epicardial biology in the adult heart and provide a basis for manipulating epicardial states in a new era of regenerative therapies.

## ACKNOWLEDGMENTS

The authors thank the Technology Center for Genomics & Bioinformatics for their single-cell library construction and sequencing services.

## SOURCES OF FUNDING

This work was supported by National Institutes of Health (NIH) grants T32-AR065972, T32-HL069766 (to D. Wong), and R01-HL172852 (to P. Quijada). Additional support was provided by the American Heart Association (AHA) Career Development Award (19CDA34590003 to P. Quijada), the Integrative Biology and Physiology at University of California, Los Angeles (UCLA) Eureka Fellowship (to D. Wong), the Asrican Sophie & Jack Award (to D. Wong and J. Cheng), and the UCLA Broad Stem Cell Research Center Innovation Award (to P. Quijada).

## Notes

### Competing Interest Statement

The authors have declared no competing interest.

