## Supplemental Material for "Single-Cell Profiling of Dynamic Epicardial Cell States During Myocardial Infarction"

**The Supplemental Material file includes the following:**

**Supplemental Methods**
**Supplemental Figures 1-13**
**Supplemental Tables 1 and 2**
**Supplemental References**

### Supplemental Methods

#### Data Availability

Detailed descriptions of all experimental procedures and materials are available in the Supplemental Material. Single-cell RNA sequencing data have been deposited in the National Center for Biotechnology Information Gene Expression Omnibus (GEO) under accession #GSE336312. Fetal epicardial cell single-cell RNA-sequencing can be found under the GEO accession #GSE154715.

#### Sham / Myocardial Infarction Surgery

Mice were anesthetized by exposure to a 2% isoflurane/oxygen mixture and were administered Buprenorphine (2.5mg/kg) subcutaneously. A midline cervical incision was made to expose the trachea for intubation with a PE90 plastic catheter. The catheter was connected to a Harvard minivent supplying supplemental oxygen, with a tidal volume of 225-250 µL and a respiratory rate of 130 strokes/min (Harvard Apparatus Model 845). Surgical plane anesthesia was subsequently maintained at 1-1.5% isoflurane. The skin was incised, and the chest cavity was opened at the 4th intercostal space. The mouse was placed on a heating pad (half-inch plexiglass between the animal and the heating pad). Oral intubation was performed by placing PE 90 tubing into the mouth and advancing it slowly into the trachea. Mechanical P.I. ventilation began with a tidal volume of approximately 0.4 ml at 130 breaths/min. Maintenance anesthesia was maintained at 1.25-1.75% isoflurane. After intubation, a midline incision was made between the sternum and the left internal artery. Alternatively, a lateral incision was made in the fourth intercostal space. The heart was exposed, and the left coronary artery was ligated intramurally 2mm from the origin with an 8-0 proline suture. Lungs were reinflated, and the chest was closed in two layers; the ribs (inner layer) were closed with 6-0 coated Vicryl sutures in an interrupted pattern. The skin was closed using 6-0 nylon or silk sutures in a subcuticular manner. Anesthesia was discontinued, and once the mouse was breathing independently, it was removed from the ventilator and allowed to recover in a clean cage on a heated pad. Sham surgeries were performed identically to MI procedures without an attempt at ligation of the left anterior artery and mice in this group were used as controls.

**Adult mouse epicardial cell isolation for single-cell RNA sequencing (scRNA-seq)**

8-week-old male and female *Wt1*$^{CreERT2/+}$*; R26*$^{tdTomato/+}$*; PDGFR*$\alpha^{nGFP/+}$ mice were given an intraperitoneal injection of tamoxifen (10 mg/kg) for five consecutive days. After a week-long washout period, mice were subject to sham or myocardial infarction surgery. Mice were sacrificed on 7 or 14 days post-MI or sham surgery following 7 days through the following: anesthetized with an intraperitoneal injection of 0.4 mL of ketamine-xylazine cocktail (13 mg/mL ketamine and 0.88 mg/mL xylazine in PBS) and sacrificed by cervical dislocation. The thoracic cavity was opened, and the hearts were removed and placed in a petri dish containing PBS. The heart was squeezed gently with a tweezer to remove blood from the chambers, and the atria, vessels, and external connective tissue were removed. Left ventricles were microdissected, minced into 1-2 mm pieces with scissors, and placed in scintillation vials containing a magnetic stir bar. Enzyme mix from the Neonatal Heart Dissociation Kit (Miltenyi Biotec, 130-098-373) was added according to the manufacturer's instructions to the scintillation vials, and the hearts were incubated at 37℃ for 45 minutes on a stir plate. Digestion was stopped by adding cold, sterile 0.5% bovine serum albumin dissolved in PBS. The digested heart solutions were filtered through a 70μm strainer, centrifuged, and the cell pellet was resuspended in PBS containing 0.04% BSA and maintained on ice before sorting. For each condition/time point, cells were collected from Sham (n=5), 7dMI (n=5), and 14dMI (n=2) mice and processed into pooled libraries. Sample size per surgery group was based on the consistency in infarct size across animals at the same time-point, and mice were excluded from heart digestion and single-cell collection if they did not exhibit at least a 40% reduction in ejection fraction following MI surgery. Sham mice needed to show a healthy ejection fraction of at least 60%; due to consideration of several variables, such as post-surgical mortality and infarct size, a priori sample size selection was not used. Randomization was completed to allocate animals to non-injured or injured groups; however, because isolation days were specific, we did not randomize groups or perform blinding due to pooling animals for each time-point. Due to the scheduling of cell sorting, surgeries were completed on various days; however, the surgeon, housing location, surgery room, and downstream machines used for cell sorting were maintained consistently throughout the experiment.

**Fluorescence-activated cell Sorting (FACS)**

Single-cell suspensions were FACS-purified to enrich for epicardial lineage cells prior to scRNA-seq. Cells were stained in PBS/0.04% BSA on ice and filtered immediately before sorting. Dead cells were excluded using a viability dye (DRAQ5). Doublets were excluded by sequential FSC-A/FSC-H and SSC-A/SSC-H gating. Epicardial-lineage cells were identified as tdTomato$^{+}$ (*Wt1*-lineage traced), and GFP fluorescence was recorded to capture fibroblast-associated epicardia/EPDC-like states. Sorted cells were collected into chilled tubes containing PBS/0.04% BSA and immediately processed for 10X Genomics library preparation.

**RNA isolation, cDNA synthesis, and Real-Time Quantitative Reverse Transcription PCR**

To perform gene expression analyses, isolated epicardial cells were sorted directly into TRIzol. RNA was extracted from cells according to the manufacturer's protocol. Extracted RNA was quantified using a NanoDrop Spectrophotometer (Thermo Fisher Scientific) and then converted into complementary DNA (cDNA) using Verso cDNA Synthesis Kit (Thermo Fisher Scientific, AB1453A) following the manufacturer's instructions. RT-qPCR reactions were prepared with SsoAdvanced Universal SYBR Green Supermix (Bio-Rad, 1725274), and specific primers designed for each target gene are listed in **Supplemental Table 1**. The reactions were run on a CFX Opus Real-Time PCR (Bio-Rad). Data analysis was performed using the ΔΔCt method, and gene expression levels were normalized to *18s* for mice.

| Supplemental Table 1. Primer Sequences for RT-qPCR | | | |
|---|---|---|---|
| **Species** | **Gene Target** | **Forward Primer** | **Reverse Primer** |
| *mus musculus* | 18S | CATGGCCTCAGTTCCGAAAA | CGAGCCGCCTGGATACC |
| *mus musculus* | COL1A1 | TAGGCCATTGTGTATGCAGC | ACATGTTCAGCTTTGTGGACC |
| *mus musculus* | MSLN | TGGTGAGGTCACATTCCACT | TGGACAAGACCTACCCACAA |
| *mus musculus* | PDGFRA | GGGAGAGAAACAAACGGAGGA | GCTCCTGAGACCTTCTCCTTCTA |
| *mus musculus* | POSTN | AAGCTGCGGCAAGACAAG | TCAAATCTGCAGCTTCAAGG |
| *mus musculus* | TCF21 | CATTCACCCAGTCAACCTGA | CCACTTCCTTCAGGTCATTCTC |
| *mus musculus* | WT1 | ATCCGCAACCAAGGATACAG | GGTCCTCGTGTTTGAAGGAA |

**scRNA-seq library construction and sequencing**

Sorted cells were processed using the 10X Genomics Chromium platform (Next GEM Single Cell 3' v3.1). cDNA amplification and library construction were performed according to the manufacturer's protocol. Qubit-quantified libraries and fragment size distributions were assessed using Bioanalyzer/TapeStation before sequencing. Sequencing was performed on an Illumina platform with paired-end 2x100 bp read configuration.

**scRNA-seq Data Preprocessing and Quality Control**

Base calling and demultiplexing were performed using Illumina software, and FASTQ files were aligned to the mouse reference genome (GRCm38). Cell Ranger v 8.0.0 was used to generate gene-by-cell count matrices. A custom reference including the tdTomato and eGFP transgenes was used for alignment. Cells were retained if they expressed >200 detected genes and had >500 UMI counts, with upper cutoffs applied to remove potential multiplets (nFeature_RNA < 5000, nCount_RNA < 50,000). Cells with high mitochondrial transcript content were excluded (percent.mt < 25%). Unbiased clustering using the Seurat R package, with visualization via uniform manifold approximation and projection (UMAP) dimension reduction, was performed to identify cells with distinct lineage identities and transcriptional profiles[1].

**Normalization, integration, and batch correction**

For adult epicardial cell datasets (Sham, 7dMI, 14dMI), normalization and variance stabilization were performed using SCTransform (Seurat). Datasets were integrated using Seurat's anchor-based workflow (SelectIntegrationFeatures, PrepSCTIntegration, FindIntegrationAnchors, IntegrateData). For fetal-adult comparisons, embryonic epicardial scRNA-seq datasets (E12.5 and E16.5) were integrated with adult datasets using the same SCTransform + anchor-based integration workflow to enable joint clustering and cross-condition state comparisons.

**Cell Clustering**

Principal component analysis (PCA) was performed on the scaled (SCT) expression matrix using highly variable genes, and the top PCs were used to construct a shared nearest-neighbor (SNN) graph (FindNeighbors). Clustering was performed using Louvain/Leiden community detection (FindClusters) across a range of resolutions, and the selected resolution was guided by cluster stability, marker-gene specificity, and biological interpretability. Cluster annotation was based on canonical markers and differential expression signatures, including epicardial/mesothelial markers (*Wt1*, *Msln*, *Krt19*, etc), fibroblast/activated stromal markers (*Postn, Pdgfra, Fibronectin*, etc), endothelial markers (*Pecam1, Kdr)*, immune markers (*Ptprc*), and cardiomyocyte markers (*Tnnt2*), enabling identification and exclusion of non-epicardial contaminants where appropriate.

**Differential Gene Expression Analysis and Pathway Enrichment**

Cluster marker genes were identified using Seurat (FindAllMarkers / FindMarkers) with the Wilcoxon rank-sum test, requiring expression in at least 10% of cells (min.pct) and a minimum log-fold change threshold of 0.25. P-values were adjusted for multiple testing using Bonferroni or Benjamini-Hochberg false discovery rate (FDR) as indicated. Condition-specific differential expression (7dMI vs Sham; 14dMI vs 7dMI) was performed within matched cell states/clusters to isolate injury-responsive programs while controlling for cell identity. Pathway enrichment analyses were performed using clusterProfiler (enrichGO for Gene Ontology Biological Process). Enrichment significance was assessed using hypergeometric tests with FDR correction (Benjamini-Hochberg).

#### Cell-cell communication and ligand-receptor inference

To infer changes in intercellular signaling after MI, we applied CellChat[2] to integrated Seurat objects. Communication probability was computed, aggregated at the pathway level, and compared at the 7dMI condition to identify signaling pathways active after MI. Sender/receiver roles were defined using interaction-strength metrics, and key pathways were prioritized based on biological relevance.

#### Immunofluorescence and in situ hybridization assays

Whole hearts harvested following sham or MI surgeries following arrest of the heart in diastole by perfusing hearts with 0.1 M Cadmium Chloride (Sigma-Aldrich, 655198)/Potassium Chloride (Sigma-Aldrich, P3911), followed by perfusion with PBS for 5 minutes, then 10% neutral buffered formalin for 10 minutes at 80–100 mmHg via retrograde cannulation of the abdominal aorta. Retroperfused hearts were removed from the thoracic cavity and placed in 10% neutral buffered formalin for 16-24 hours before submission for processing. Hearts were perfused, harvested, and fixed in 10% neutral buffered formalin for 18-24 Hours at room temperature on a rocking platform. After fixation, tissue was dehydrated in an ethanol series, then xylene, before being embedded in paraffin wax and cut into 5 µm sections using a microtome. After sectioning, slides were allowed to dry overnight at room temperature and stored with desiccants for long-term storage. We used the RNAscope Multiplex Fluorescent V2 Assay (Advanced Cell Diagnostics, 323100) for in situ hybridization on formalin-fixed, paraffin-embedded (FFPE) tissue per the manufacturer's instructions. Slides were mounted with ProLong Gold Antifade Mountant (Thermo Fisher Scientific, P36934) before being imaged on Nikon AX R NSPARC. RNA probes and secondary detection methods are listed in **Supplemental Table 2.** Image acquisition parameters were held constant within experiments.

| **Supplemental Table 2. RNAscope mRNA Probes for Fluorescence-based In Situ Hybridization** | | |
|---|---|---|
| **Gene Target** | **RNAscope Catalog #** | **Akoya Biosciences # for Fluorescent Detection and Dilution** |
| MSLN | 443241-C2 | Akoya Biosciences OPAL 570 (FP1488001KT); 1:750 |
| POSTN | 418581-C3 | Akoya Biosciences OPAL 520 (FP1487001KT); 1:750 |
| TDTOMATO | 317041 | Akoya Biosciences OPAL 690 (FP1497001KT); 1:750 |

**Statistics**

Data were presented as mean +/− standard error of the mean (SEM) for bar graph data. Statistical analyses were performed using unpaired two-tailed Student's t-test when comparing two groups and One-Way and Two-Way ANOVA when comparing multiple groups. Tukey post-test was used to correct multiple comparisons made by One-Way ANOVA, and Sidak post-test was used to correct multiple comparisons made by Two-Way ANOVA. All measurements in this paper were acquired from different biological samples, and no samples were measured repeatedly. Bar graph data analysis, Graph Pad Prism 11 for macOS. A p-value less than 0.05 was considered statistically significant.

**Code availability**

All analyses were performed using standard protocols with previously described R packages. The R scripts are available upon request.

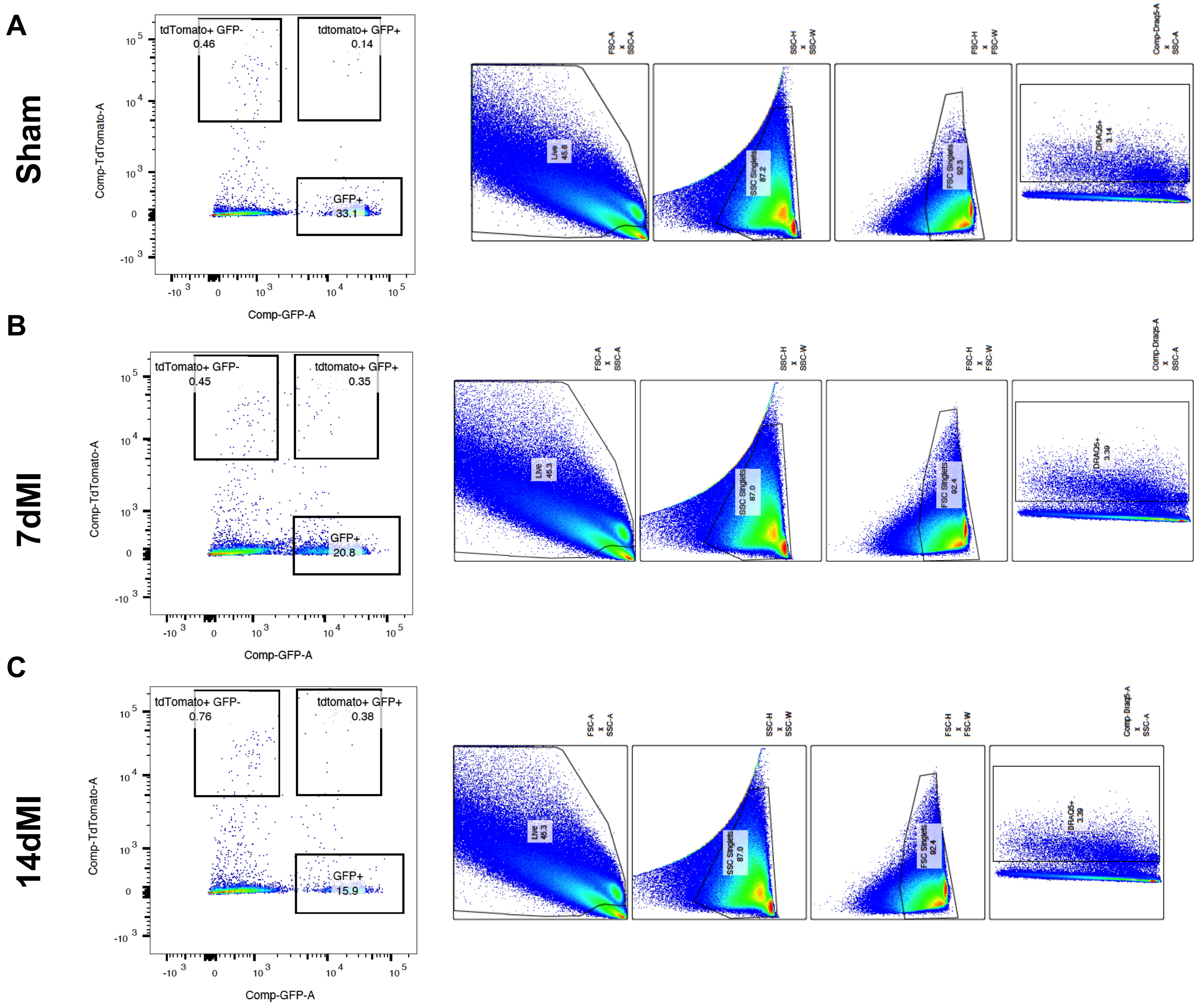


**Supplemental Figure 1. FACS-based gating strategy for the isolation of epicardial-derived cells following sham or myocardial infarction surgery.**

**(A-C)** Fluorescence-activated cell sorting (FACS) plots representing the gating workflow used to isolate viable/nucleated (DRAQ5$^+$) single epicardial-derived (*Wt1*-lineage) cells for scRNA-seq following sham surgery or myocardial infarction. Left panels show fluorescence profiles of *Wt1*$^{CreERT2/+}$ *; R26*$^{tdTomato/+}$ *; Pdgfra*$^{nGFP/+}$ hearts, gated on tdTomato (epicardial lineage) and GFP (Pdgfra, fibroblast marker). Right panels represent forward/side scattering and fluorescent plots to exclude doublet and non-nuncleated cells (DRAQ5$^-$). Data is representative of one animal per condition.

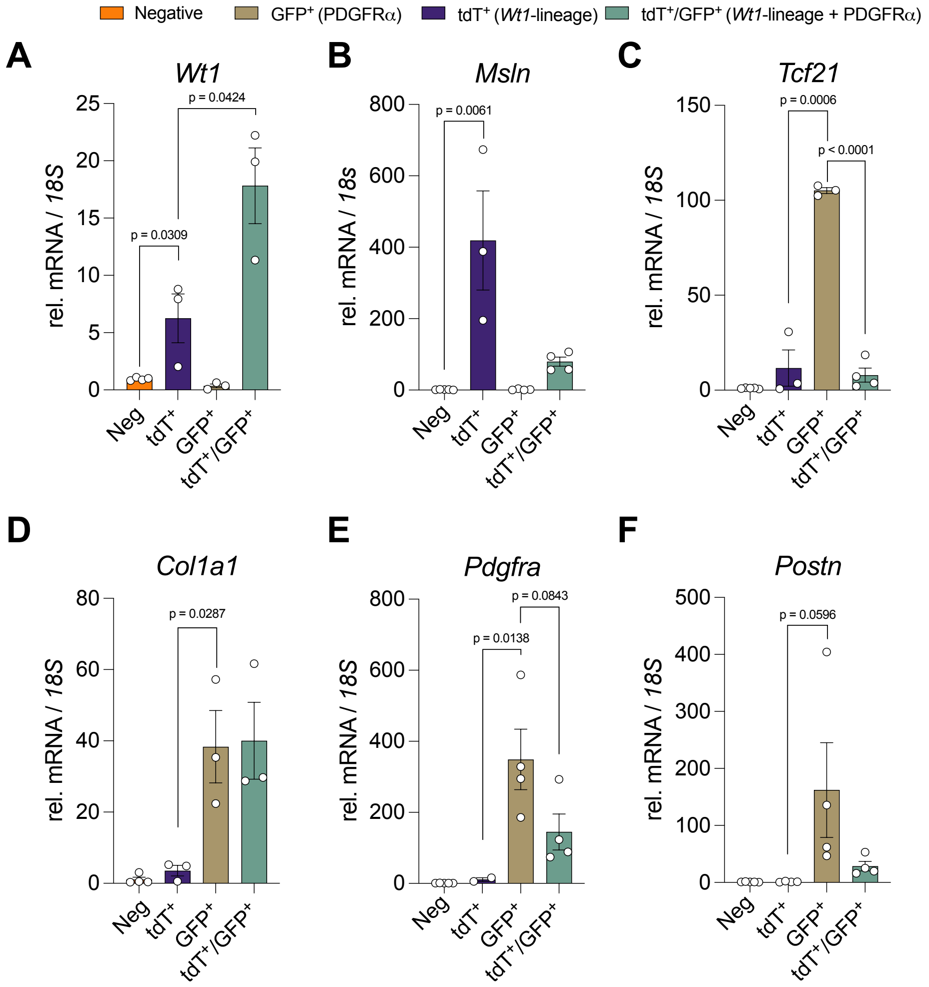


**Supplemental Figure 2. Expression of epicardial and and fibroblast gene in epicardial subpopulations via FACS.**
**(A-B)** Expression of epicardial cell markers *Wt1* and *Msln.*
**(C-F)** Expression of fibroblast markers T*cf21, Col1a1, Pdgfra, and Postn.*
Data represented as mean +/- SEM; signficance determined by unpaired two-tailed Student's t-test. Data is presented with biological replicates: N=3-4 of the negative, tdTomato+, GFP+ and tdTomato+/GFP+ fractions isolated via FACS following 7 days of myocardial infarction.

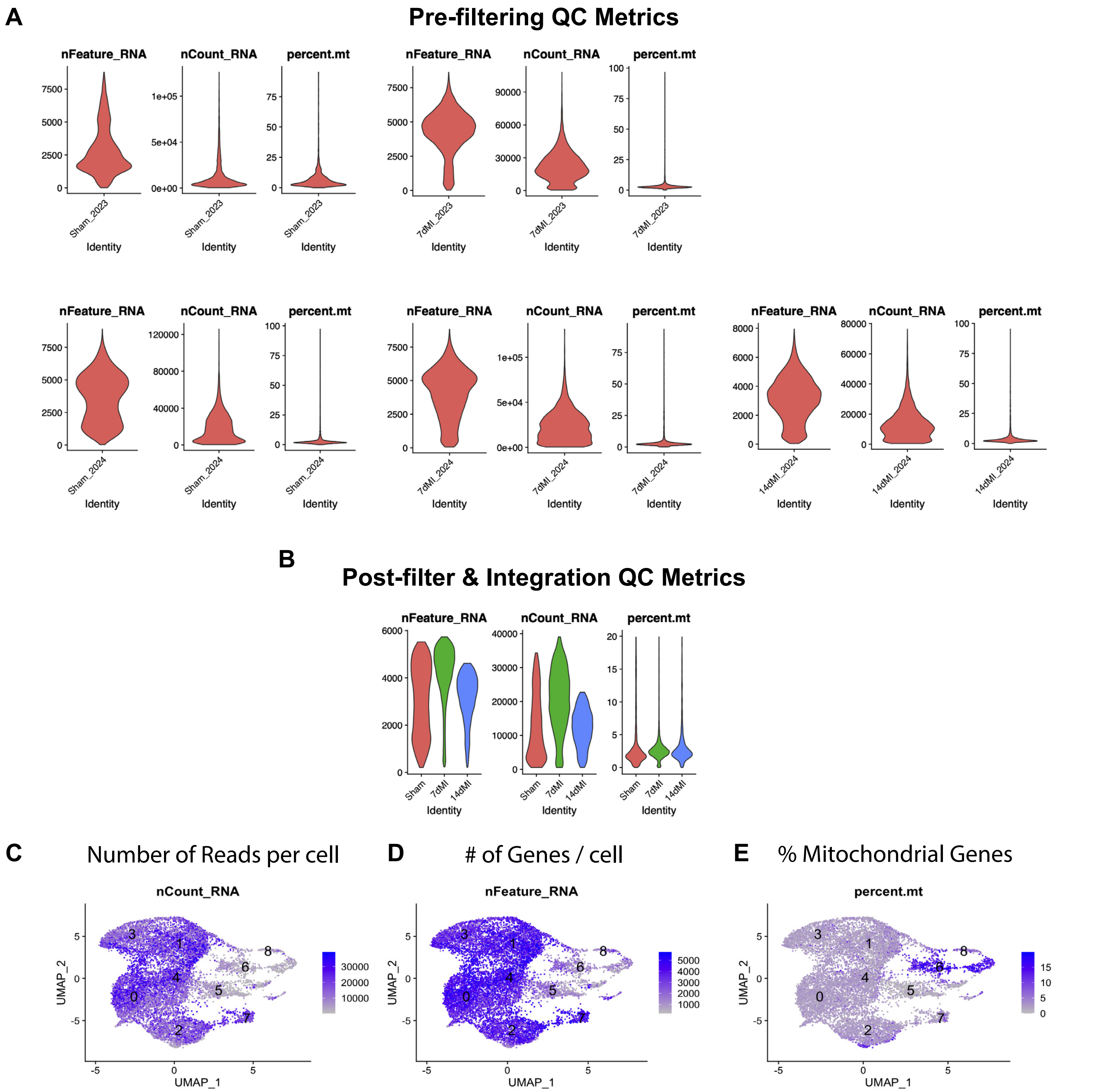


**Supplemental Figure 3. Single-cell RNA sequencing quality control and filtering of epicardial cells. (A)** Pre-filtering quality control (QC) metrics for all samples, shown as violin plots of detected genes per cell (nFeature_RNA), number of reads per cell (nCount_RNA), and percentage of mitochondrial transcripts (percent.mt). **(B)** Post-filter and integration QC metrics across all conditions (Sham, 7dMI, 14dMI). **(C-E)** UMAP visualizations of QC metrics across the integrated dataset: **(C)** number of reads per cell, **(D)** number of detected genes per cell, and **(E)** percentage of mitochondrial transcripts.
Data are presented as biological replicates: N=5 for Sham, N=5 for 7dMI, and N=2 for 14dMI.

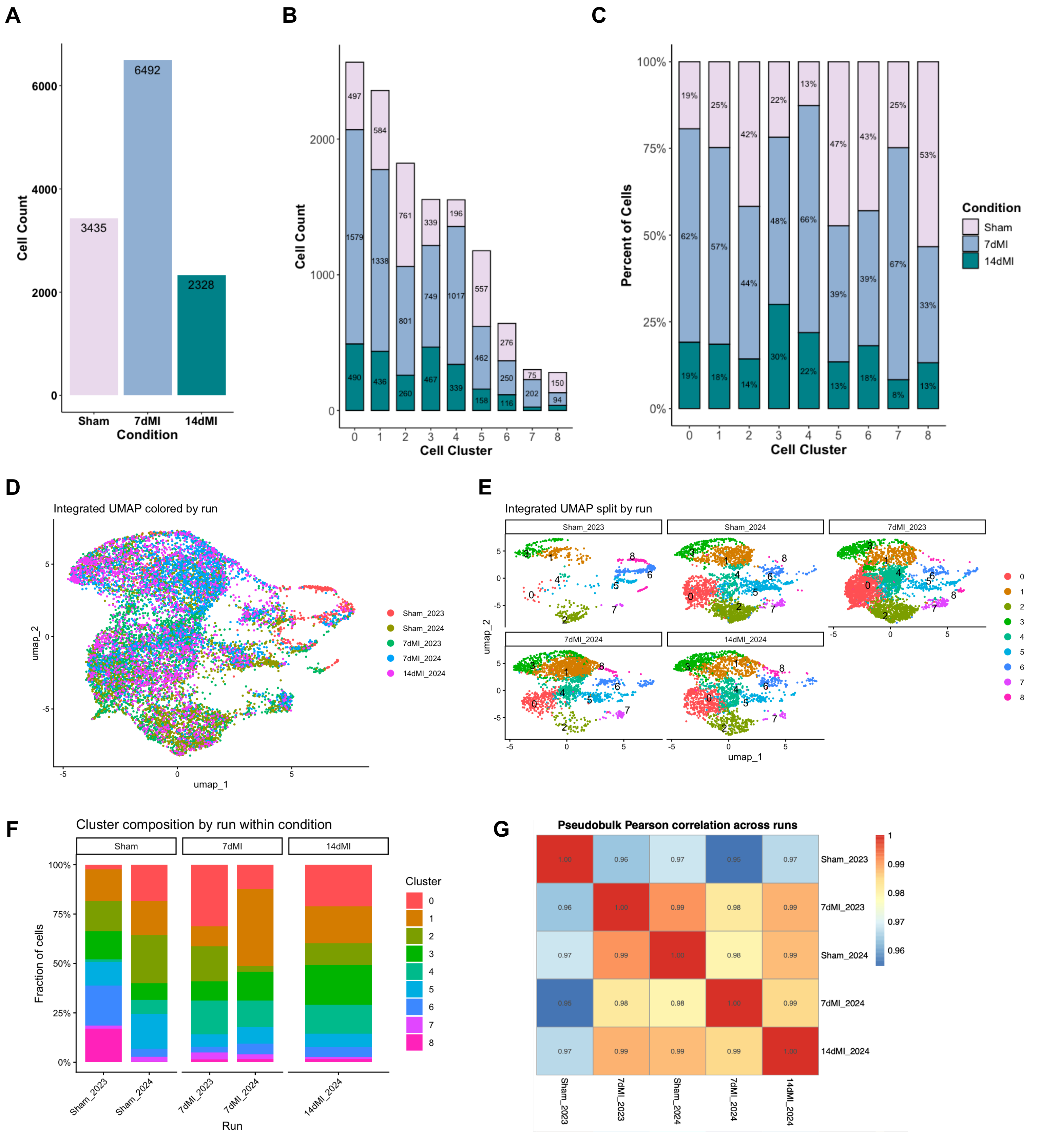


**Supplemental Figure 4. Assessment of batch correction and run-level reproducibility across scRNA-seq datasets.**
**(A)** Total tdTomato⁺ epicardial-lineage cells captured from sham, 7dMI, and 14dMI conditions following quality control filtering.
**(B)** Distribution of total cell counts across all clusters, separated by condition.
**(C)** Relative percentage of cells from each condition across all clusters.
**(D)** UMAP of integrated epicardial lineage cells colored by sequencing run after SCTransform-based Seurat Integration.
**(E)** UMAP split by run and colored by cluster identity, showing contribution of each run to the same major clusters.
**(F)** Cluster composition by run within condition, plotted as the fraction of cells assigned to each cluster.
**(G)** Pseudobulk Pearson correlation heatmap across runs generated from raw RNA counts aggregated by run, demonstrating high overall transcriptomic similarity and no obvious outlier run.
Data are presented as biological replicates: N=5 for Sham, N=5 for 7dMI, and N=2 for 14dMI.

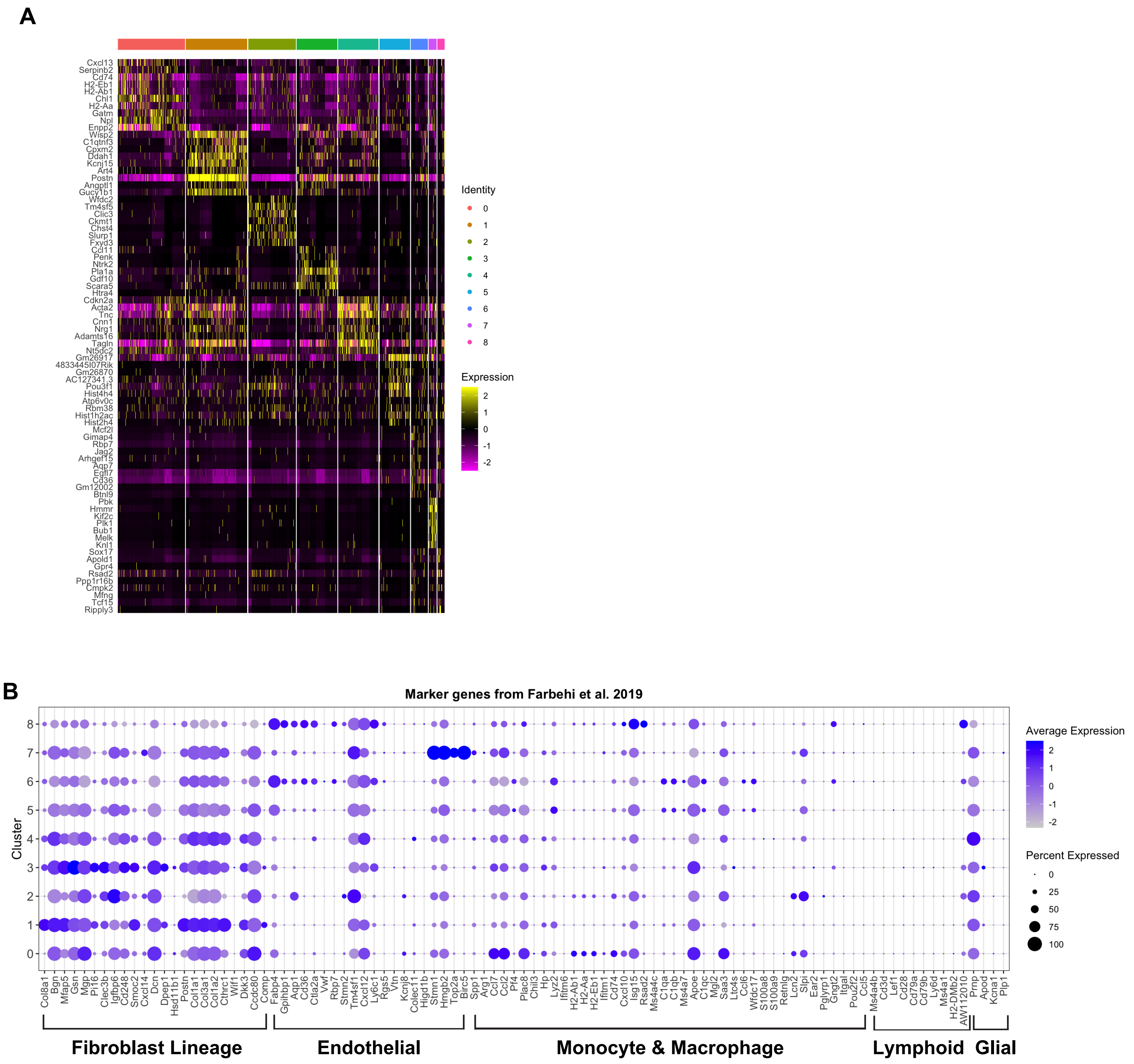


**Supplemental Figure 5. Epicardial cell cluster identification.**
**(A)** Heat map showing the expression of the top 10 most differentially expressed genes in epicardial cell clusters. Scale represents relative expression values.
**(B)** Dot plot showing expression of published marker genes from Farbehi et al. (2019) across epicardial cell clusters. Genes are grouped by lineage-associated programs to define fibroblasts, endothelial cells, monocytes and macrophages, lymphoid cells, and glial cells.
Data are presented as biological replicates: N=5 for Sham, N=5 for 7dMI, and N=2 for 14dMI.

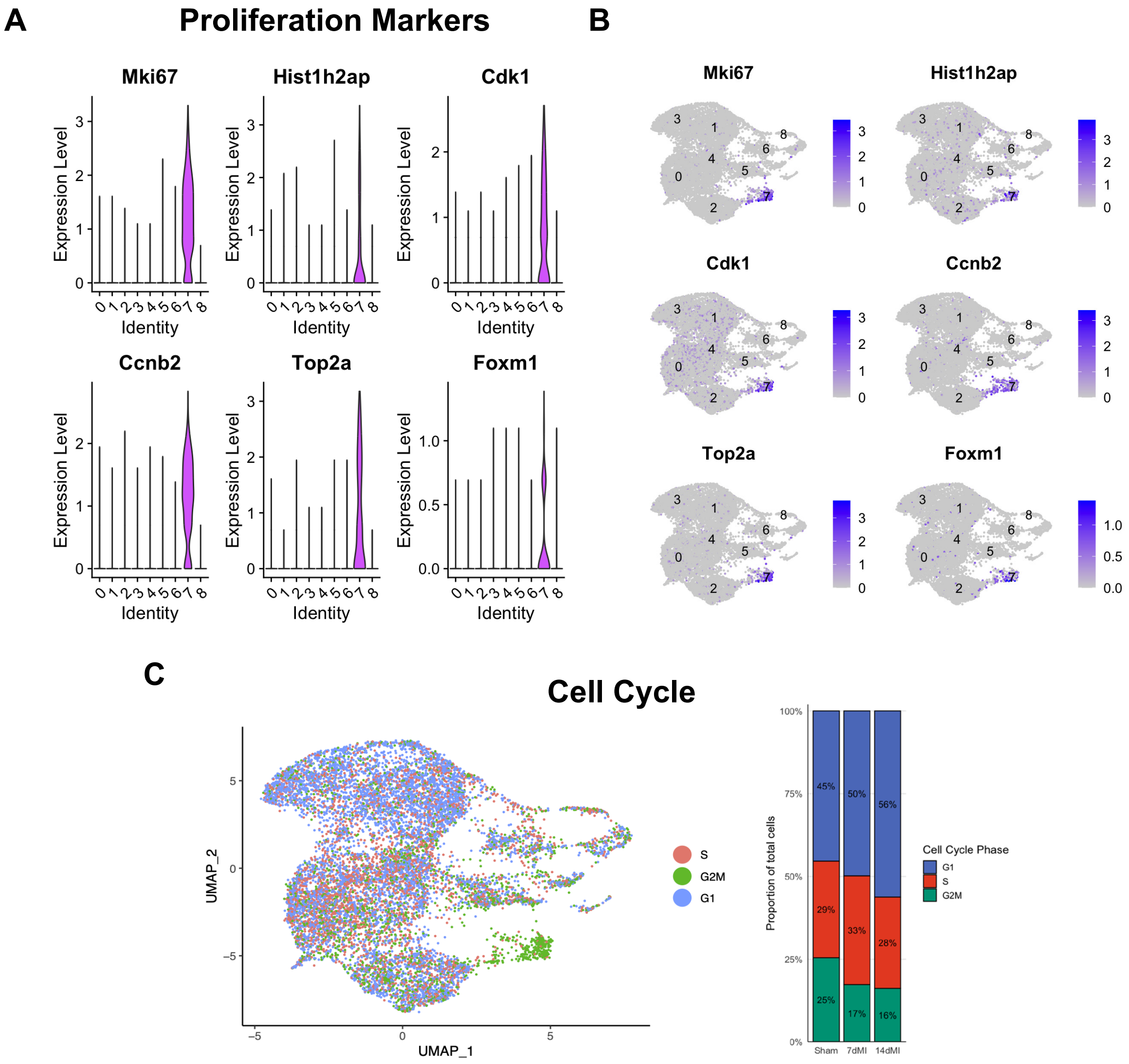


**Supplemental Figure 6. Assessment of cell cycle activity in epicardial cells.**
**(A)** ViolinPlots showing expression of key cell-cycle associated genes across all clusters.
**(B)** FeaturePlots showing spatial distribution of cell-cycle gene markers in **(A)**.
**(C)** Distribution of cells in S, G2/M, or G1 phase based on cell cycle scoring.
Data are presented as biological replicates: N=5 for Sham, N=5 for 7dMI, and N=2 for 14dMI.

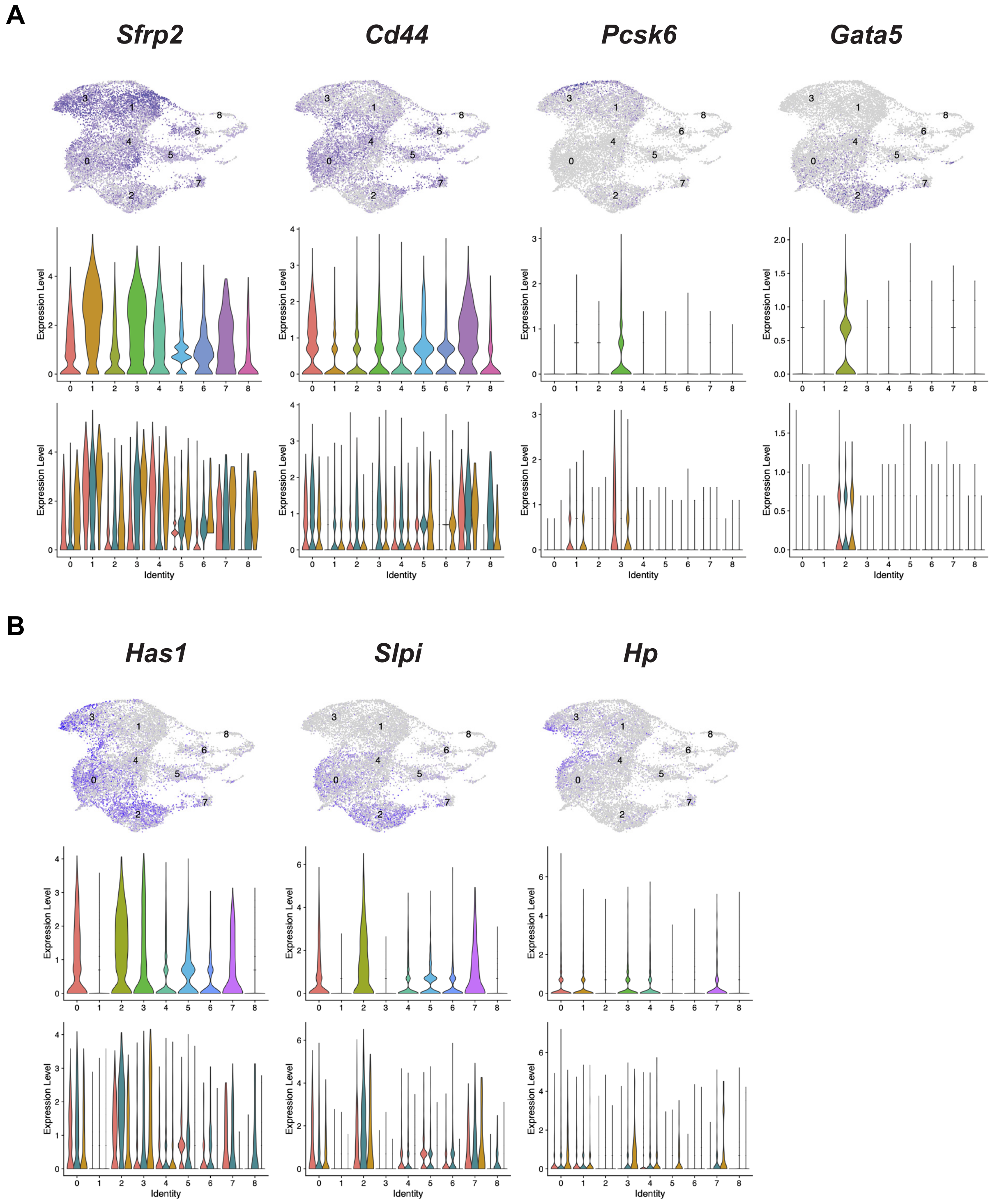


**Supplemental Figure 7. Expression of epicardial marker genes in uninjured and infarcted hearts.**
**(A-B)** Top row*:* FeaturePlots, violin plots by cluster (middle row), and violin plots grouped by condition (bottom row; Sham, 7 days post-MI, 14 days post-MI). Data are presented as biological replicates: N=5 for Sham, N=5 for 7dMI, and N=2 for 14dMI.
**(A)** Expression of epicardial markers previously identified by Hesse et al., 2021.
**(B)** Expression of adult human epicardium-specific markers identified by Knight-Schrijver et al., 2022.

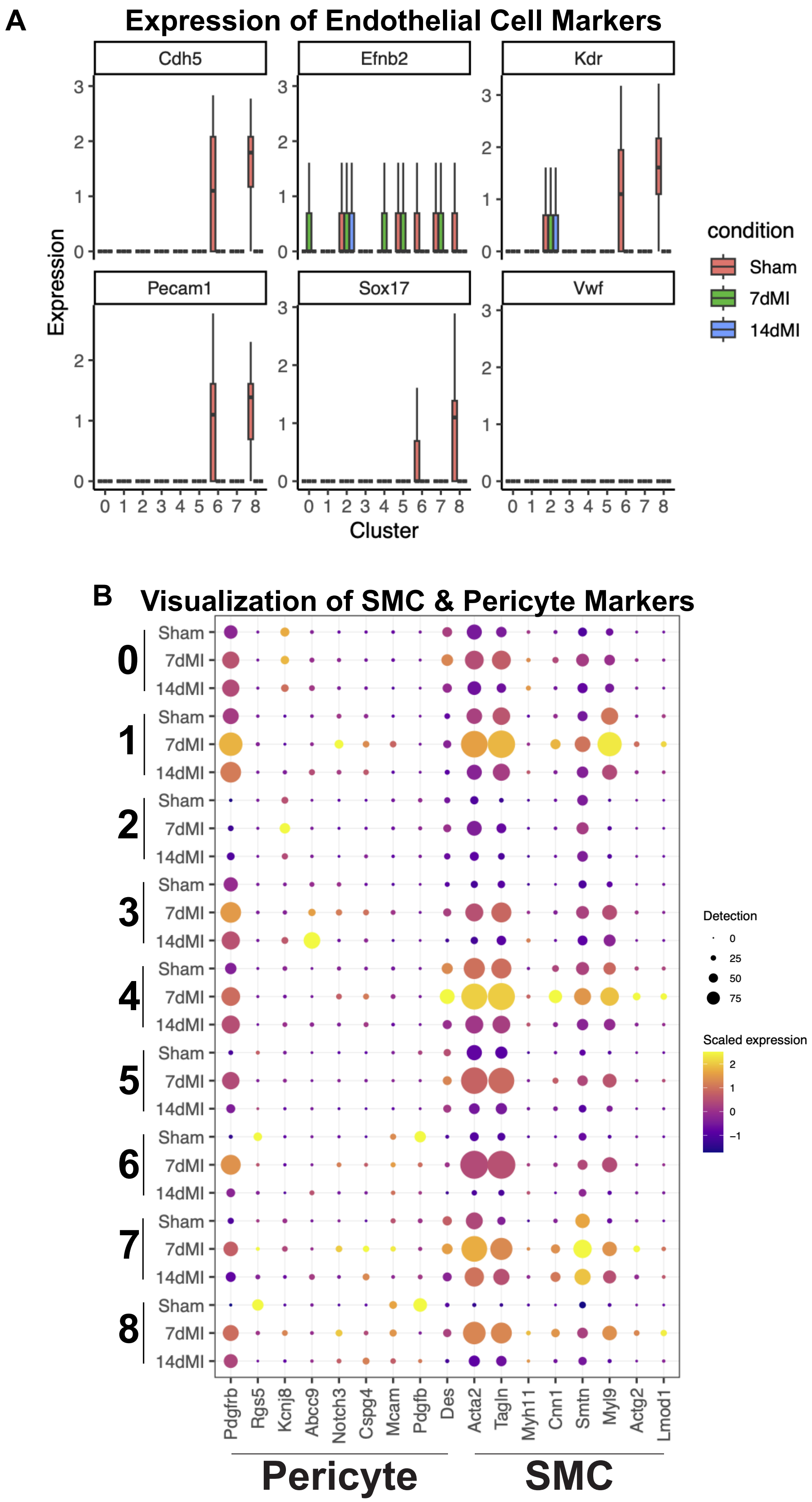


**Supplemental Figure 8. Expression of endothelial cell, smooth muscle cell, and pericyte marker genes.**
**(A)** Boxplots showing expression of canonical endothelial cell markers across clusters and condtions.
**(B)** Dot plot displaying the expression of pericyte and smooth muscle cell associated genes across clusters and conditions. Dot size reflects the gene, and color indicates average scaled expression.
Data are presented as biological replicates: N=5 for Sham, N=5 for 7dMI, and N=2 for 14dMI.

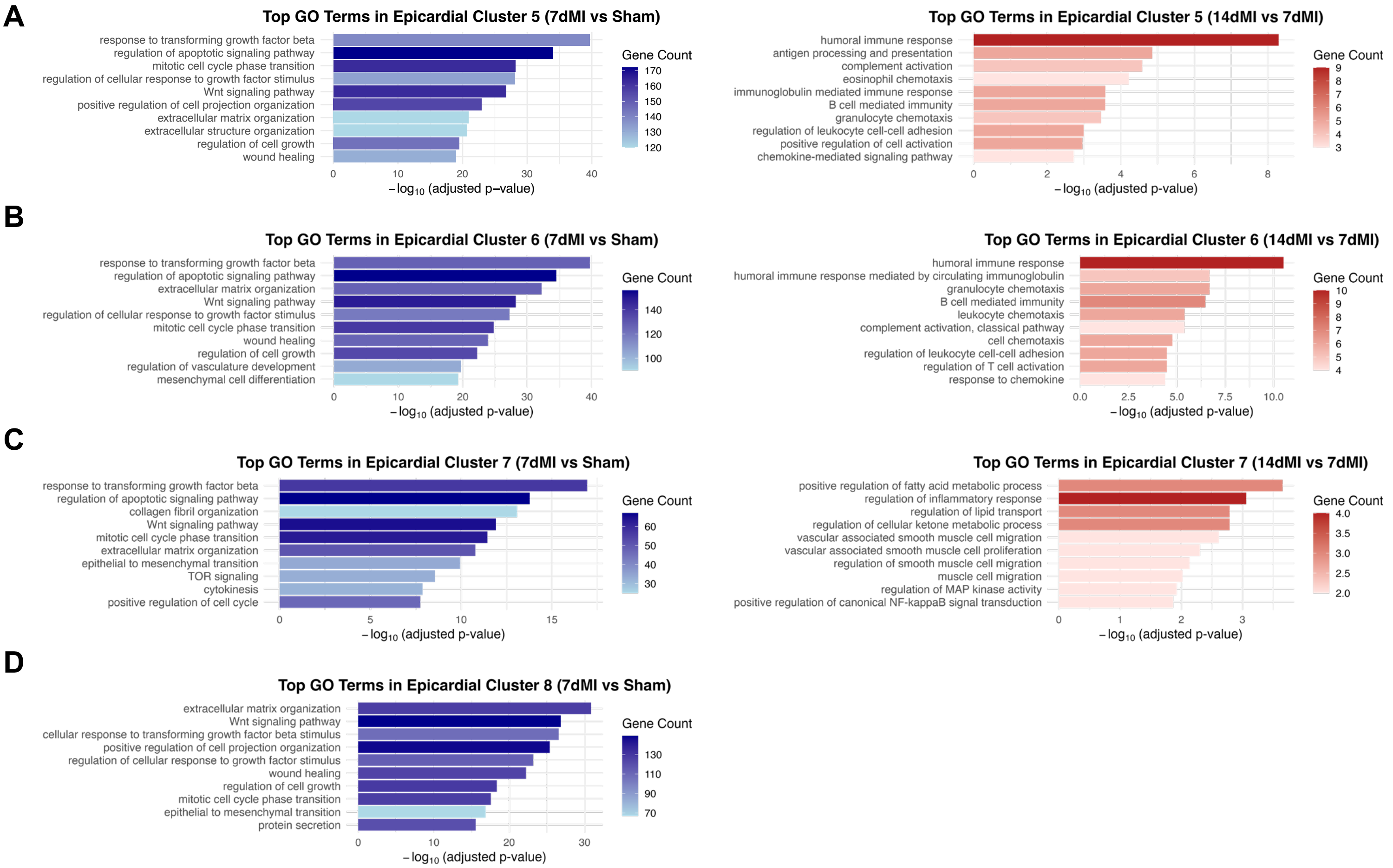


**E**

| Cluster | 7dMI vs Sham | | 14dMI vs 7dMI | |
|---|---|---|---|---|
| | Upregulated | Downregulated | Upregulated | Downregulated |
| 0 | 4325 | 47 | 78 | 5275 |
| 1 | 6318 | 66 | 132 | 4868 |
| 2 | 4885 | 77 | 48 | 3830 |
| 3 | 5366 | 91 | 131 | 4153 |
| 4 | 3681 | 71 | 105 | 4542 |
| 5 | 4383 | 60 | 29 | 3359 |
| 6 | 3633 | 159 | 27 | 2698 |
| 7 | 1470 | 7 | 8 | 353 |
| 8 | 3747 | 173 | 4 | 794 |
| | min.pct = 0.1<br>logfc.threshold = 0.25 | | min.pct = 0.05<br>logfc.threshold = 0.1 | |

**Supplemental Figure 9. Cluster-specific Gene Ontology (GO) enrichment analysis across conditions.**
**(A-D)** Bar plots ofrepresentative GO Biological Process terms in epicardial clusters 5-8 comparing 7dMI vs. Sham (left, blue) and 14dMI vs 7dMI (right red). Bars represent the -log(adusted p-value) of each GO term, with gene count indicated by color intensity. **(E)** Summary table showing the number of upregulated and downregulated differentially expressed genes (DEGs) per cluster in each comparison. DEG thresholds were set at min.pct = 0.1 and log2FC > 0.25 for 7dMI vs Sham, and min.pct = 0.05 and log2FC > 0.1 for 14dMI vs 7dMI.
Data are presented as biological replicates: N=5 for Sham, N=5 for 7dMI, and N=2 for 14dMI.

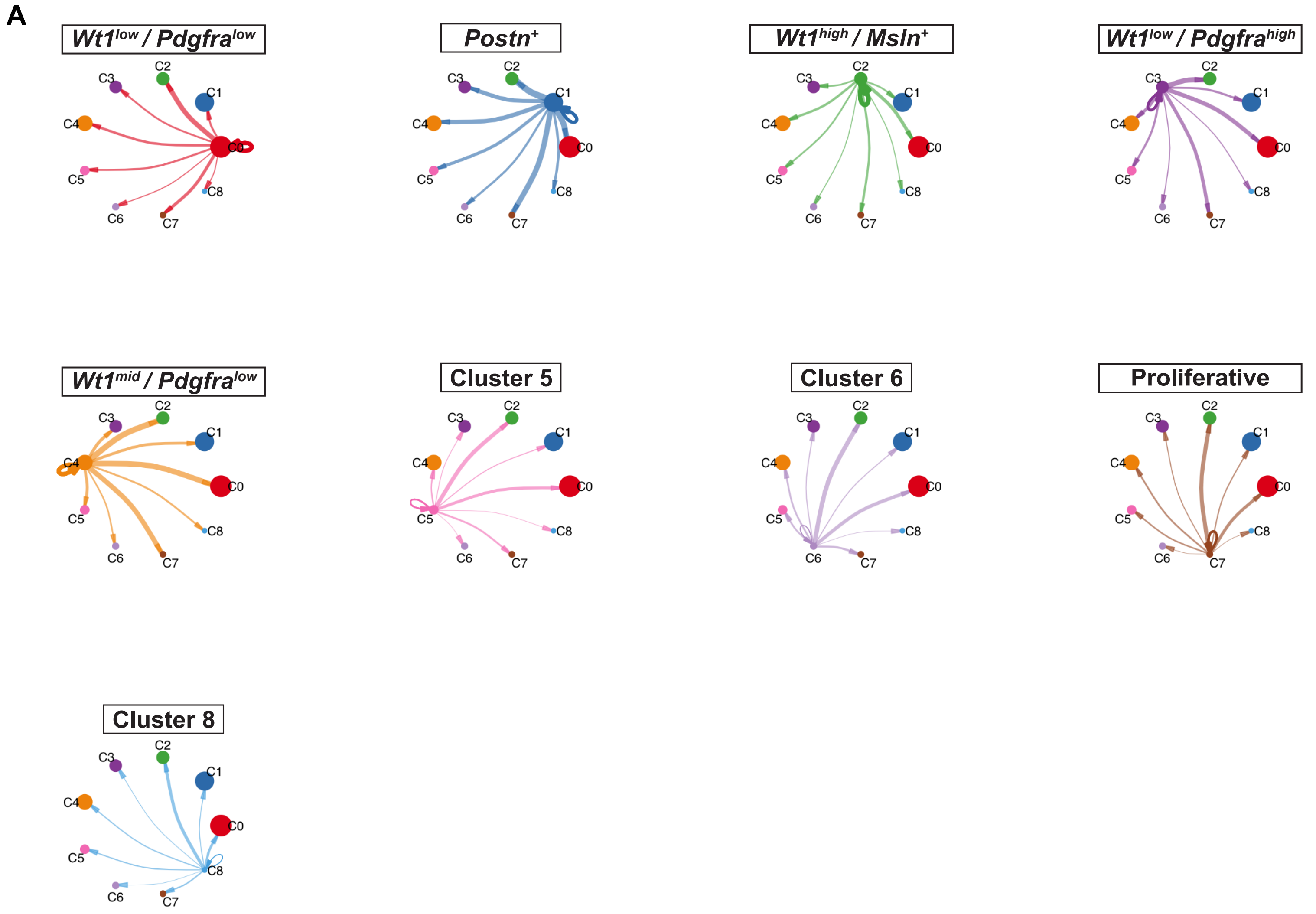


**Supplemental Figure 10. Cluster-specific cell-cell signaling inferred by CellChat.**
**(A)** Circle plots showing outgoing signaling from each epicardial cells cluster. Labeled node represents the designated sender cluster, and all nodesare sized according to the relative number of cells in each cluster. Edge/arrow thickness is proportional to the communication probability (interaction strength) across all pathways and self-referencing arrows represent autocrine signaling. Data are presented as biological replicates: N=5 for Sham, N=5 for 7dMI, and N=2 for 14dMI.

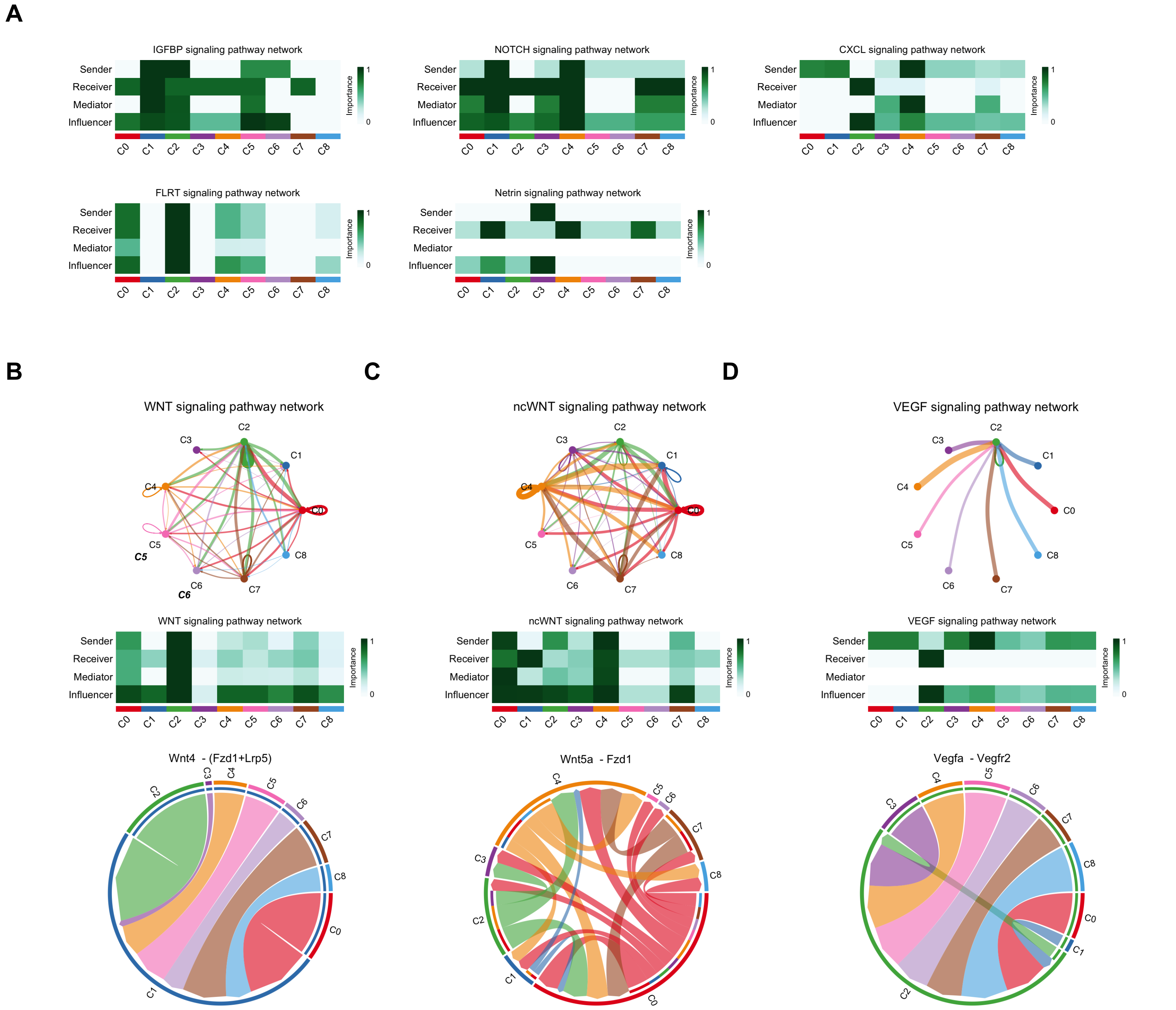


**Supplemental Figure 11. CellChat analysis of epicardial cell-cell communication and dominant ligand-receptor interactions at 7 days post-MI.**
**(A)** Signaling pathway heatmaps from CellChat displaying the relative importance of each epicardial cluster (C0-C8) as Sender, Receiver, Mediator, or Influencer for selected pathways (IGFBP, NOTCH, CXCL, FLRT, Netrin).
**(B-D)** Pathway focused summaries for WNT **(B)**, non-canonical WNT (ncWNT) **(C)**, and VEGF **(D)** signaling. Top: aggregated cell-cell communication networks between epicardial clusters, with directed edges representing inferred signaling flow (sender --> receiver) and edge thickness reflecting interaction strength. Middle: pathway-specific signaling role heatmaps. Bottom: chord diagrams highlighting the most dominant ligand-receptor interaction within each pathway.
Data are presented as biological replicates: N=5 for Sham, N=5 for 7dMI, and N=2 for 14dMI.

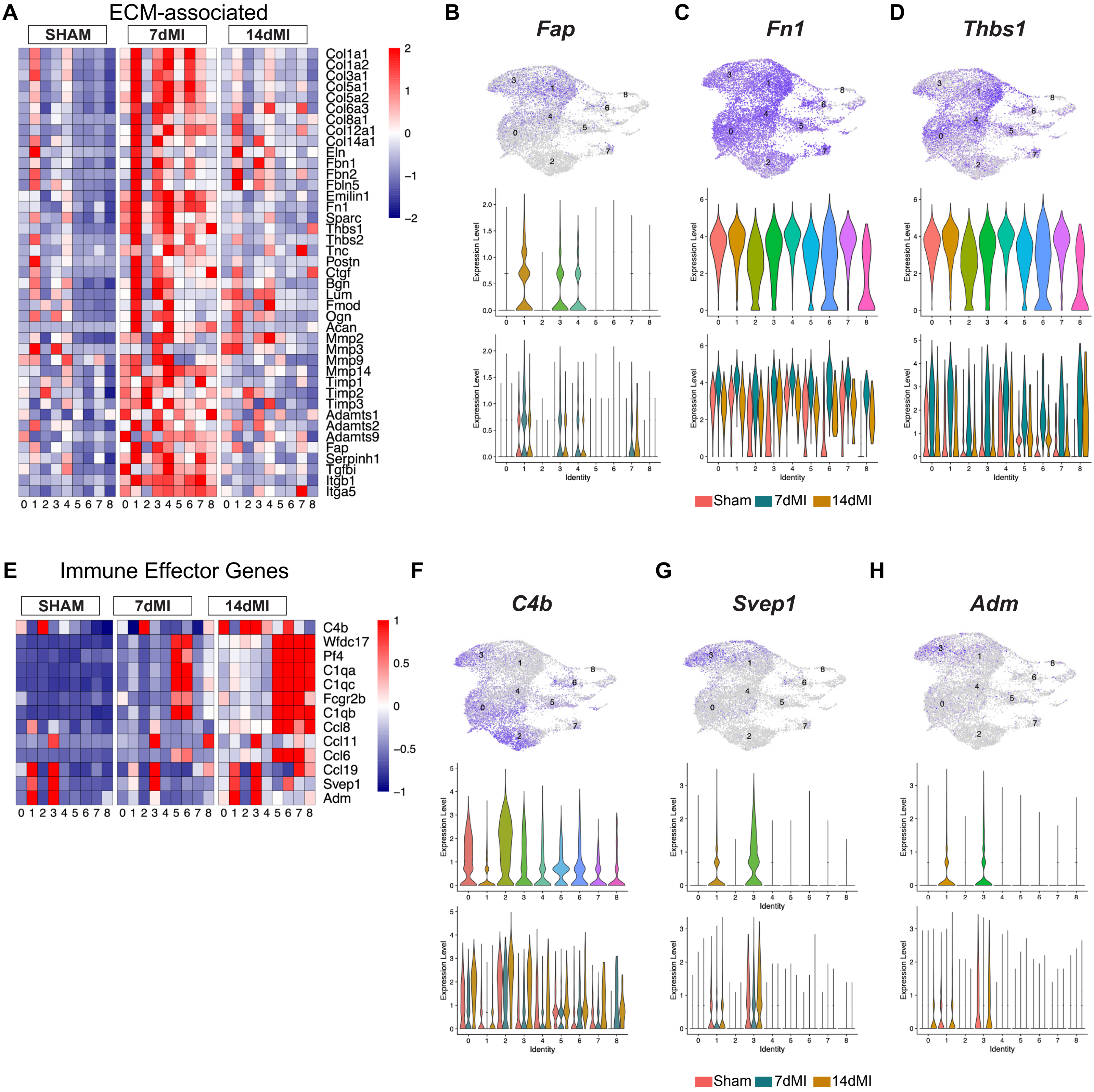


**Supplementary Figure 12. Extracellular matrix and immune response gene expression dynamics following MI.**
**(A)** Heatmap displaying Z-scored expression of extracellular matrix (ECM) genes across conditions
**(B-D)** Feature and violin plots showing cluster-specific expression of *Fap, Eln,* and *Mmp2.*
**(E)** Heatmap displaying Z-scored expression of immune effector genes across conditions.
**(F-H)** Feature and violin plots showing cluster-specific expression of *C4b, Svep,* and *Adm.*
Data are presented as biological replicates: N=5 for Sham, N=5 for 7dMI, and N=2 for 14dMI.

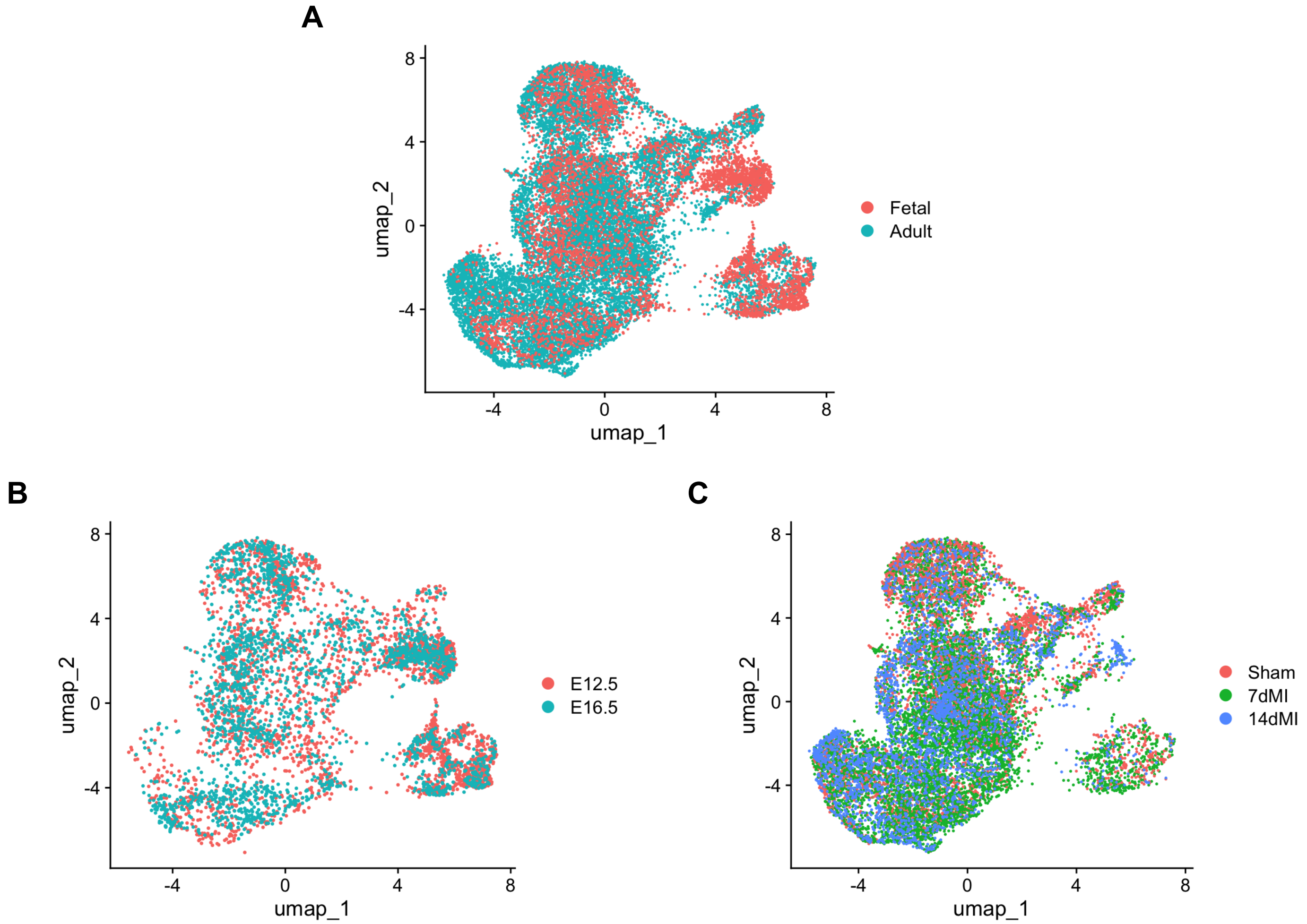


**Supplemental Figure 13. Integrated fetal and post-MI adult epicardial cells.**
**(A-C)** UMAP visualization of integrated epicardial cells from fetal (E12.5, E16.5) and adult post-MI hearts (Sham, 7dMI, 14dMI).
**(A)** UMAP colored by developmental stage group (Fetal vs. Adult)
**(B)** UMAP colored by fetal time point (E12.5 vs. E16.5)
**(C)** UMAP colored by adult injury condition (Sham, 7dMI, 14dMI)
Data are presented as biological replicates: N=5 for Sham, N=5 for 7dMI, and N=2 for 14dMI.
